# The high mutational sensitivity of *ccdA* antitoxin is linked to codon optimality

**DOI:** 10.1101/2022.01.15.476443

**Authors:** Soumyanetra Chandra, Kritika Gupta, Shruti Khare, Pehu Kohli, Aparna Asok, Sonali Vishwa Mohan, Harsha Gowda, Raghavan Varadarajan

## Abstract

Deep mutational scanning studies suggest that synonymous mutations are typically silent and that most exposed, non active-site residues are tolerant to mutations. Here we show that the *ccdA* antitoxin component of the *E.coli ccdAB* toxin-antitoxin system is unusually sensitive to mutations when studied in the operonic context. A large fraction (∼80%) of single-codon mutations, including many synonymous mutations in the *ccdA* gene shows inactive phenotype, but they retain native-like binding affinity towards cognate toxin, CcdB. Therefore, the observed phenotypic effects are largely not due to alterations in protein structure/stability, consistent with a large region of CcdA being intrinsically disordered. *E. coli* codon preference and strength of ribosome-binding associated with translation of downstream *ccdB* gene are found to be major contributors of the observed mutant phenotypes. In select cases, proteomics studies reveal altered ratios of CcdA:CcdB protein levels *in vivo*, suggesting that the *ccdA* mutations likely alter relative translation efficiencies of the two genes in the operon. We extend these results by studying single-site synonymous mutations that lead to loss of function phenotypes in the *relBE* operon upon introduction of rarer codons. Thus, in their operonic context, genes are likely to be more sensitive to both synonymous and non-synonymous point mutations than inferred previously.

## Introduction

Mutations that lead to changes in the protein’s amino acid sequence are often expected to alter the protein structure, stability and/or activity and are referred to as missense mutations. Unlike missense mutations, synonymous mutations alter only the DNA (and mRNA) sequence without affecting the amino acid sequence because of the degeneracy of the genetic code (Crick et al. 1961). Synonymous mutations were classically believed to be silent in their effects on protein function. While this is often true, there have been several pioneering studies that show prominent effects of synonymous mutations on protein function (Komar et al. 1999; Kimchi-Sarfaty et al. 2007; Sander et al. 2014). Synonymous mutations are also reported to be associated with over 50 human diseases (Sauna and Kimchi-Sarfaty 2011). Some of the mechanisms known to be involved in the altered phenotypes, observed upon introduction of multiple synonymous mutations, are due to the effects on protein expression and folding (Buhr et al. 2016; Rodnina 2016), transcription (Zhao et al. 2021), as well as mRNA stability due to changes in secondary structure or altered mRNA decay rates (Presnyak et al. 2015; Hanson and Coller 2018). Such effects are often found to be brought about by the bias due to unequal codon usage (Sharp et al. 1986; Karlin et al. 1998; Quax et al. 2015) and tRNA abundance (Gorochowski et al. 2015) as well as effects on translation initiation (Li et al. 2012), elongation or termination (Rodnina 2016).

Most deep mutational scanning (DMS) studies involving parallel high-throughput investigation of phenotypic fitness in large number of single-site mutants of proteins have revealed synonymous mutations to be largely neutral (Hietpas et al. 2011; Jiang et al. 2013; Melamed et al. 2013; Roscoe et al. 2013; Wu et al. 2014), with the exception of studies such as in β-lactamase where a very small number of synonymous mutants (∼2%) display < 50% of WT activity (Firnberg et al. 2016) and in *Saccharomyces cerevisiae* Hsp90 where one out of fifteen synonymous mutations tested shows significant loss of fitness (Fragata et al. 2018). Several low throughput studies have investigated phenotypic fitness effects of small sets of synonymous and non-synonymous (missense) single-nucleotide or single-codon mutants and attempted to elucidate molecular mechanisms responsible for the observed patterns of distribution of fitness effects (DFE) (Sanjuán et al. 2004; Carrasco et al. 2007; Domingo-Calap et al. 2009; Lind et al. 2010; Peris et al. 2010; Cuevas et al. 2012; Schenk et al. 2012; Bailey et al. 2014; Agashe et al. 2016; Kristofich et al. 2018; Lebeuf-Taylor et al. 2019). These studies either use adaptive evolution experiment to screen for emerging beneficial mutants or use site-directed mutagenesis (SDM) to construct a small subset of single-site mutants to study their fitness. Several single-site synonymous mutants have been identified to exhibit lower fitness than WT with reduced growth rates, while others have higher fitness with improved growth rates in various bacterial genes under selection pressure (Lind et al. 2010; Schenk et al. 2012; Bailey et al. 2014; Agashe et al. 2016; Kristofich et al. 2018). A number of studies in viruses also identify small numbers of synonymous mutants to be deleterious or even lethal (Sanjuán et al. 2004; Carrasco et al. 2007; Domingo-Calap et al. 2009; Wu et al. 2014), as analyzed and detailed in an useful review on synonymous mutants in laboratory evolution experiments (Bailey et al. 2021). However, most of these DFE studies show deleterious synonymous mutations to have smaller effects than non-synonymous mutations. A study of DFE in two ribosomal proteins reveals a large fraction of mutants to be weakly deleterious and fitness effects of synonymous and non-synonymous mutants to be comparable (Lind et al. 2010). While some of the studies cite altered expression levels (higher transcript levels) or mRNA stability alterations causing the observed synonymous mutational effects, the molecular basis of altered fitness in synonymous mutants has not been elucidated in most of the above described cases. For non-synonymous mutations, the large effects of amino acid changes on proteins structure, stability and function may often overshadow and obscure the mutational effects on mRNA transcript levels, structure, stability and translation.

Investigation of mutational effects on protein function has been studied extensively in mono-cistronic proteins or in isolated, heterologous expression systems. Studies of bacterial genomes however reveal a high occurrence of operonic systems comprising of functionally related protein coding genes proximally spaced on a DNA stretch, under a shared promoter (Jacob and Monod 1961; Yanofsky 1981; Balázsi et al. 2005; Price et al. 2006; Zhang et al. 2012). Operonic genes are typically separated by less than 20 base pairs (Moreno-Hagelsieb and Collado-Vides 2002). Operons exhibit characteristic tightly regulated mRNA expression from a common promoter, followed by translation of the gene products from the same mRNA molecule. The translation re-initiation efficiency of genes clustered in operons is often dependent on the space between genes (Chemla et al. 2020). Consequently, the DNA and mRNA sequence features are expected to play an important role in gene expression and regulation in operons. Moreover, transcription and translation are known to be spatiotemporally coupled in prokaryotes (Byrne et al. 1964; Stent 1964; Miller et al. 1970). In tightly regulated operonic systems, downstream functions are often acutely dependent on the relative levels of gene products of the operonic genes. Thus such operonic systems provide a sensitive and relatively unexplored readout to study phenotypic effects of synonymous mutations.

Bacterial type II Toxin-Antitoxin (TA) systems, where the antitoxin and the toxin proteins are expressed as part of a single operon, serve as an attractive model to study co-regulation of gene expression as well as transcriptional and translational coupling (Masachis et al. 2018). These TA systems usually code for an upstream labile antitoxin protein responsible for negating the toxicity of the downstream toxin protein, the latter can cause growth arrest or cell death (Goeders and Van Melderen 2014; Deter et al. 2017). The *E. coli* F-plasmid borne *ccd* operon encodes a TA system that comprises of a labile 72 residue CcdA antitoxin, that prevents killing of the bacterial cells by binding to the 101-residue toxin component, CcdB under the control of a single auto-regulated promoter (Miki et al. 1984; Tam and Kline 1989). Both genes are co-expressed in low amounts in F-plasmid bearing *E. coli* cells, and their expression is autoregulated at the level of transcription (Afif et al. 2001). If the cell loses F-plasmid, the labile CcdA is degraded by the ATP dependent Lon protease, releasing CcdB from the complex to act on its DNA gyrase target. Free CcdB poisons gyrase by forming a CcdB-gyrase-DNA ternary complex and induces double-stranded breaks in DNA (Bahassi et al. 1999; Van Melderen 2002). This eventually leads to cell death (Van Melderen et al. 1996). Therefore, mutations which disrupt CcdA antitoxin function or lower the CcdA:CcdB ratio *in vivo*, can lead to cell death. We therefore carried out saturation mutagenesis studies of the antitoxin *ccdA*, from the F plasmid encoded CcdAB TA system in *E.coli* to understand how point mutations, especially synonymous mutations, might affect gene function and regulation in such an operonic context.

We find that the labile antitoxin CcdA in the operonic context shows considerably higher mutational sensitivity than most proteins studied to date (Gupta and Varadarajan 2018). Several synonymous point mutations in CcdA lead to loss of antitoxic function and significant reduction in cell survivals, in a manner dependent on the *E. coli* codon preference. Our data suggests that reduction in optimality of the codons upon mutation results in decreased levels of CcdA protein, likely by altering the translation efficiency. The reduced CcdA: CcdB protein ratio *in vivo* in such mutants, results in heightened toxic effects on bacterial cells. Moreover, the low CcdA: CcdB ratio also results in upregulation of *ccd* transcription, further amplifying the effect of mutation, resulting in cell death. Improved translation initiation of the downstream toxin CcdB, upon mutations in the *ccdA* gene, especially in the ribosome binding site (RBS) of CcdB, also appears to contribute to increased toxicity in cells. The *ccd* and likely most other TA operons, are highly sensitive genetic circuits that can be used to probe effects of mutations on gene function, *in vivo*. We observe that a large fraction (∼80%) of single codon variants of the *ccdA* gene (both synonymous as well as non-synonymous mutations) results in partial or complete loss of antitoxic function, these are referred to as inactive phenotypes.

## Results

### Unexpectedly high mutational sensitivity in the *ccdA* antitoxin gene, in its native operonic context

In the *ccd* operonic context, mutations in the 72 residue antitoxin CcdA that impair its antitoxic function are expected to promote cell death (Figure 1A). To understand the consequences of mutations in *ccdA* on bacterial growth, the *ccd* operon consisting of the promoter, *ccdA* and *ccdB* genes (Figure 1B) was cloned into the pUC57 plasmid. Restriction digestion sites introduced in the construct for ease in cloning resulted in a few changes with respect to the WT sequence of the *ccdAB* operon, at certain positions in the non-coding region upstream of the CcdA gene (Supplementary Figure S1). A site-saturation mutagenesis (SSM) library of CcdA was created in this operonic construct using NNK primers (Supplementary Figure S2). The library comprised of both synonymous and non-synonymous mutants of CcdA, along with ∼10% WT population arising owing to the cloning strategy (see Methods). In order to retrieve the full *ccdA* library in its operonic context, it was cloned and constructed in an *E. coli* strain (Top10 gyr) resistant to CcdB toxicity. This Top10 gyr strain has a point mutation in DNA-gyrase, that prevents CcdB binding and toxicity (see Supplementary Methods). Following selection in an *E. coli* (Top 10) strain that is sensitive to the CcdB toxicity, the operonic *ccdA* mutant library was subjected to deep sequencing. The enrichment score of each CcdA mutant was computed from the available sequencing reads after selection (obtained from plasmid isolated from the sensitive strain) versus before selection (obtained from plasmid isolated from the resistant strain), relative to corresponding reads for WT CcdA (Supplementary Figure S2 and Methods). The ES_seq_ score is defined as the relative ratio of fraction of mutant reads obtained after selection to that obtained before selection, normalized to WT reads. Normalization to WT reads allows facile estimation of the enrichment of the individual mutants with respect to WT. Low enrichment is indicative of reduced survival of a mutant with respect to WT. The ES_seq_ score can therefore be used as a measure of the relative activity of mutants, which in turn is related to but not quantitatively identical to their fitness, as discussed below.

**Figure 1.**
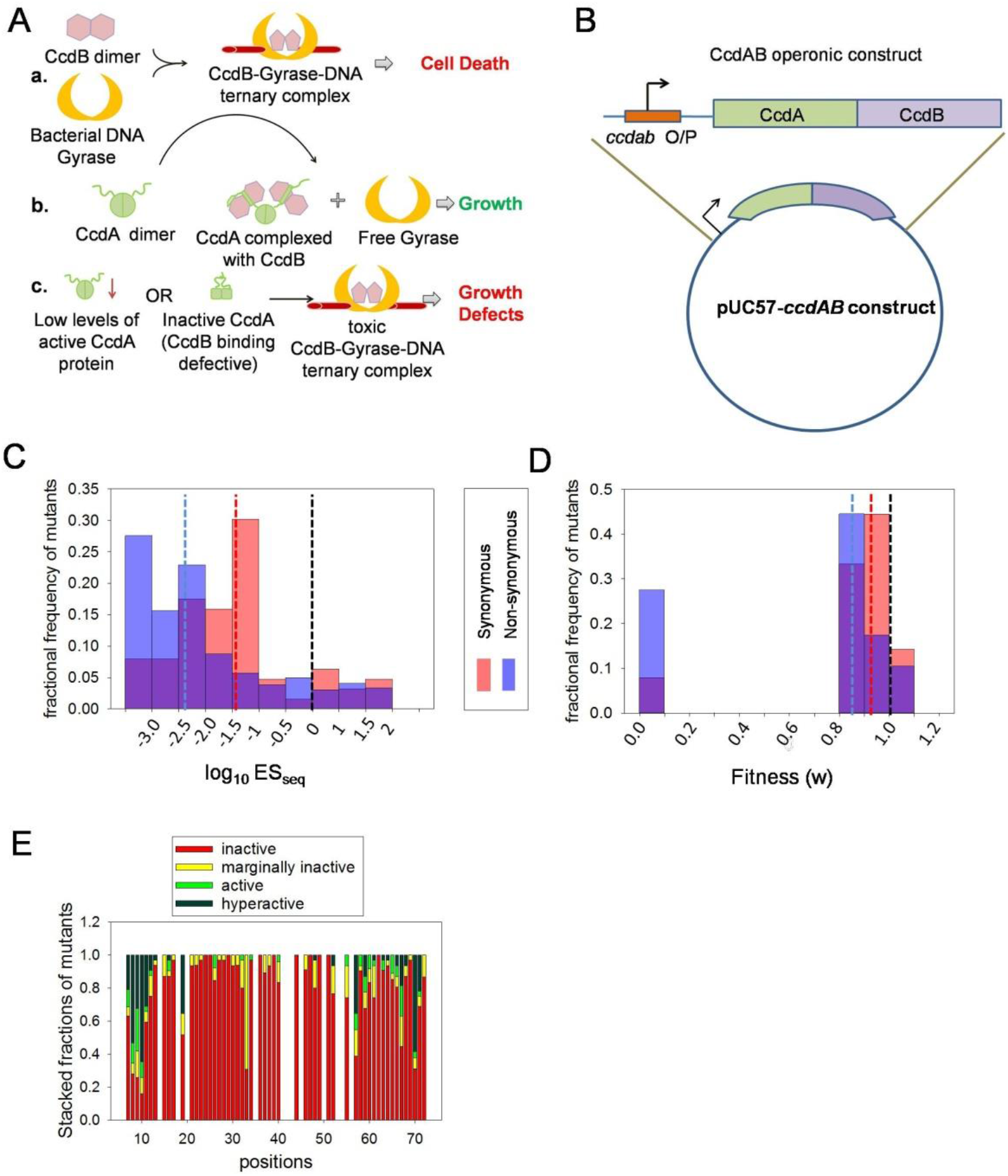
High Mutational Sensitivity of *ccdA* antitoxin gene in its operonic context in *E.coli*. (A) A schematic overview of the CcdAB TA module and the molecular mechanisms by which CcdB toxin triggers growth arrest in cells (top), how antitoxin CcdA rescues growth arrest by binding and sequestering CcdB (middle) and how mutations in CcdA can affect the antitoxic functions in the CcdAB TA system and promote CcdB mediated cell toxicity (bottom). (B) Schematic representation of *ccdAB* operon cloned in pUC57 vector. (C) Distribution of the mutational effects for synonymous (red) and non-synonymous (blue) single-site *ccdA* mutants, in the operonic context. Enrichment Score (ES_seq_) values for the *ccdA* mutants in the library are inferred through deep sequencing. A low ES_seq_ value is indicative of low antitoxic activity of the corresponding *ccdA* mutant, relative to WT (ES_seq_ =1, log_10_ES_seq_ = 0). The median values for synonymous and non-synonymous mutant subsets are shown in red and blue dashed lines respectively. WT ES_seq_ value is shown in black dashed line. The fractional frequency of mutants = number of mutants with a particular range of ES_seq_ / total number of mutants. (D) Distribution of the fitness for synonymous (red) and non-synonymous (blue) single-site *ccdA* mutants. The WT fitness and median values for synonymous and non-synonymous mutant subsets are shown in black, red and blue dashed lines respectively. (E) Phenotypic landscape for different residues in CcdA inferred from deep sequencing data. For each residue position in CcdA, the fraction of mutants showing inactive (ES_seq_≤ 0.1), marginally active (0.1<ES_seq_< 0.7), active (0.7≤ES_seq_≤ 1.5) and hyperactive phenotypes (ES_seq_> 1.5) is plotted. Only positions which have data for more than 10 mutants available have been plotted here. A large number of mutants across the length of *ccdA* display a loss of a function phenotype.

Relative fitness of mutants of microbial genes is routinely estimated from their competitive growth rates (Lind et al. 2010; Zwart et al. 2018; Lebeuf-Taylor et al. 2019) or from the change in ratio of sequencing reads of mutant versus WT over time (Hietpas et al. 2011; Roscoe et al. 2013; Fragata et al. 2018; Flynn et al. 2020). Laboratory evolution experiments of variant populations over several generations are unsuitable for accurate estimation of fitness effects in deleterious and lethal mutants. In an all-against-all competition over long timescales, mutants with higher growth rates tend to enrich exponentially over time and wipe out the slow growing or growth–defective variants completely from the population. This is beneficial in adaptive evolution experiments where the aim is to screen for improved variants or new function. However, in studies that aim to identify and investigate the fitness effects of both deleterious and beneficial variants, growth rate based screening can be disadvantageous. Unlike many previous studies that have investigated DFE of non-essential protein coding genes (Lind et al. 2010; Bailey et al. 2014; Lebeuf-Taylor et al. 2019), CcdA antitoxic function is essential for survival of cells containing the CcdAB operon. We observed that a large number of *ccdA* mutants that showed an inactive phenotype in our plate-based screen, failed to grow in liquid culture when propagated from colonies, indicating a near complete loss-of-function. This prevents estimation of relative fitness effects of *ccdA* mutants from a conventional growth based assay in liquid culture. Therefore we designed a phenotypic screen of the *ccdA* library that relies on quantifying mutants in terms of the colonies that appear after transformation. Plasmid pools used for deep sequencing were directly purified from colonies scraped from the plates.

The deep sequencing was performed in three biological replicates, which showed a high correlation (Pearson Correlation Coefficient = 0.98 approximately, Supplementary Figure S3) amongst themselves. The enrichment score (ES_seq_) was calculated and assigned only to mutants having a minimum of twenty reads in the resistant strain, for a given replicate. The enrichment scores averaged over three replicates were used to classify mutants based on their activity. The WT CcdA has an ES_seq_ value of 1. Of 2272 possible NNK single codon CcdA mutations (71 positions * 32 codons), scores could be assigned to 1528 mutants. A drastic reduction in cellular growth in the sensitive strain (after selection samples) indicated a substantial prevalence of mutants exhibiting an inactive phenotype in the library. We observed large fractions of both synonymous and non-synonymous single site mutants in *ccdA* to reduce antitoxin activity in the operonic context (Fig 1C). CcdA mutants with lower enrichment are expected to have decreased antitoxic activity, thus failing to rescue CcdB mediated cell death. The low incidence of active mutations demonstrates the extreme sensitivity of the *ccdAB* operonic construct to mutations in *ccdA*. We classified mutants having ES_seq_ values ≤ 0.1 as inactive. Surprisingly, ∼80% of all CcdA mutants have ES_seq_ values ≤ 0.1 thus displaying an inactive phenotype (Figure 1C-E). A phenotypic assay was conducted for a subset of CcdA mutants by individual transformation in resistant (control) and sensitive (selection experiment) strains, and dilution plating of the single mutants. While the estimated colony forming units (CFU) for all constructs were similar to that of WT in the resistant strain, the CFU/mL varied over a range of 10^6^ across mutants in the sensitive strain and showed good agreement with the deep sequencing derived mutational scores (Supplementary Figure S4, Table 1). Phenotypic assays of individual single mutants with ES_seq_ values less than 0.1, confirmed that they show significant growth defects (significantly lower numbers of colonies after transformation) in the sensitive strain. Even when a few colonies are obtained for these inactive mutants, these fail to grow upon inoculation into liquid media or re-streaking on agar plates. We found equal fractions (∼80%) of both synonymous (n=62) and non-synonymous (n=1466) *ccdA* mutants to be highly inactive with ES_seq_ values ≤ 0.1. The distribution of ES_seq_ scores for *ccdA* synonymous mutants (median value of 0.03) is typically higher than that for non-synonymous mutants (median value of 0.004) (Fig 1C). To compare our results with previous investigations of fitness distributions of synonymous versus non-synonymous mutants (Bailey et al. 2021), we have converted ES_seq_ scores to fitness (w) (see Methods), using a previously described formula (Bailey et al. 2014; Lebeuf-Taylor et al. 2019). The median fitness values for synonymous and non-synonymous *ccdA* mutants are 0.9 and 0.86 respectively (Figure 1D) and lower than the WT fitness of 1. In contrast to the previous investigations of mutational effects on microbial growth rate in liquid culture, the current study measures the growth of mutants in terms of colonies appearing on plates after transformation. As discussed above, several *ccdA* mutants with ES_seq_ < 0.1 and corresponding w < 0.93 fail to grow in liquid culture. On the other hand, all previously studied microbial gene variants with reported fitness values could be successfully grown in liquid culture for several generations (Lind et al. 2010; Lebeuf-Taylor et al. 2019; Bailey et al. 2021) implying that the identified inactive *ccdA* mutants are more deleterious than these previously documented cases of fitness defects. Therefore the fitness (w) measure in the current study provides an upper limit of fitness estimations in *ccdA* mutants and should not be directly compared to those in previous studies.

**Table 1.**
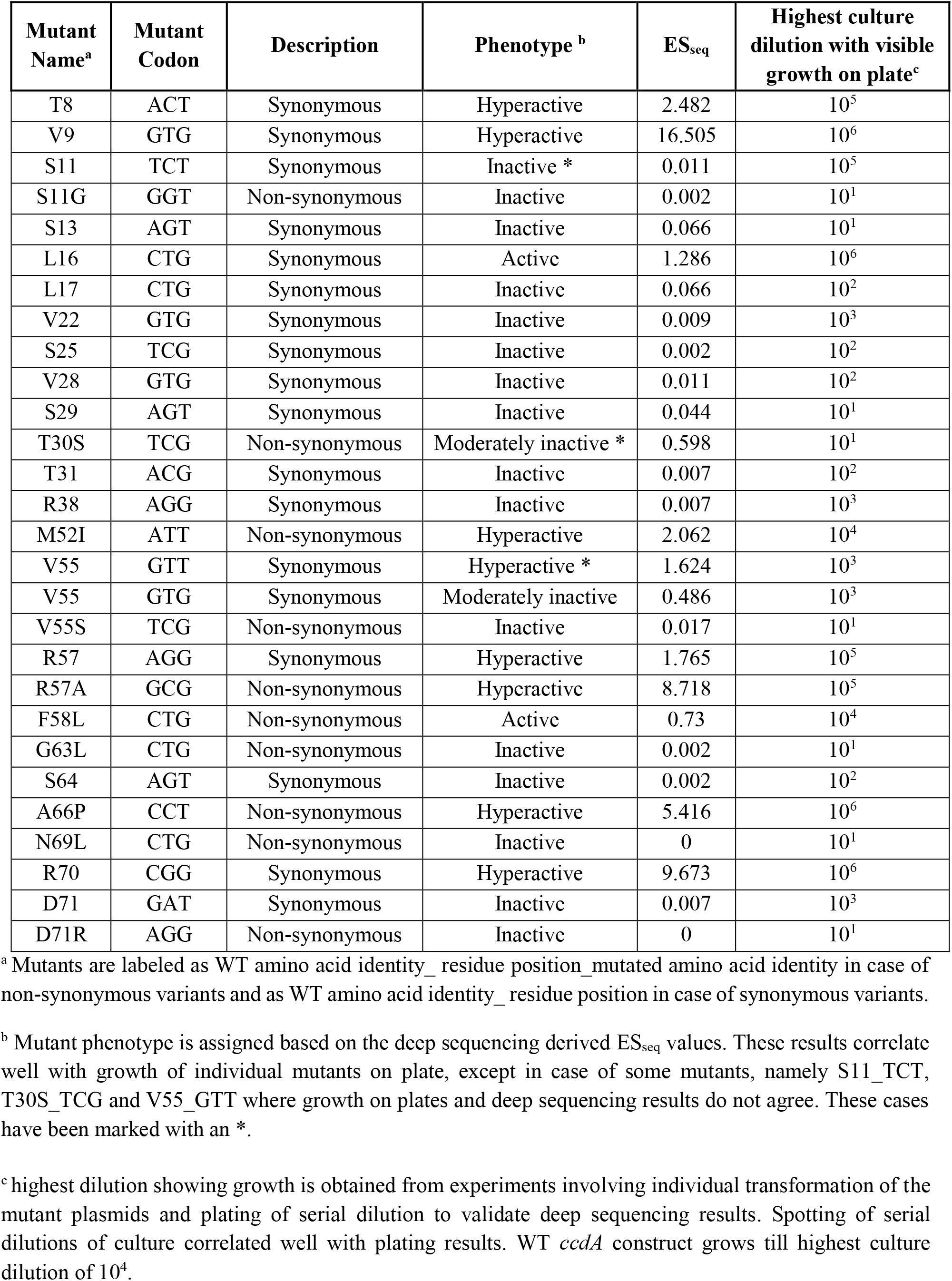
Mutant phenotypes inferred from deep sequencing and individual transformation.

### Inactive synonymous mutants and codon specific effects in *ccdAB* operonic construct

The majority of the *ccdA* single-site synonymous mutants show severe growth defects (Figure 2A and Table 2). The growth defects are inferred in terms of the number of colonies formed (and thus deep sequencing reads) by each mutant on plates following transformation in sensitive Top10 *E.coli* cells and overnight growth at 37°C (see Methods). These synonymous inactive mutants are dispersed throughout the length of the CcdA protein, with mutations at only a few specific positions at the N- and C-terminal regions showing ES_seq_ scores greater than 1. We could confidently assign ES_seq_ scores to only 62 synonymous mutants having adequate representation in the library in terms of the numbers of reads. Since we used NNK (K= G/T) primers during mutagenesis and library preparation steps, we only have 50% of all 64 codons at each position represented in the library. These 62 variants bear single codon mutations at 36 positions out of the 72 residues in CcdA. 49 of these 62 available synonymous mutants, have ES_seq_ scores less than 0.1 and can therefore be classified as highly inactive. To our knowledge such a high fraction of growth-defective single-site synonymous variants has not been previously documented in any deep mutational scan or site-directed mutagenesis study. The central stretch of CcdA (residues 20-55) was especially sensitive to synonymous mutations, with all documented synonymous mutations in this region being highly inactive. Synonymous mutations at the N terminal and C-terminal regions of CcdA resulted in both active and inactive variants. We also observed that most synonymous active mutants were hyperactive in comparison to WT with ES_seq_ > 1.5.

**Figure 2.**
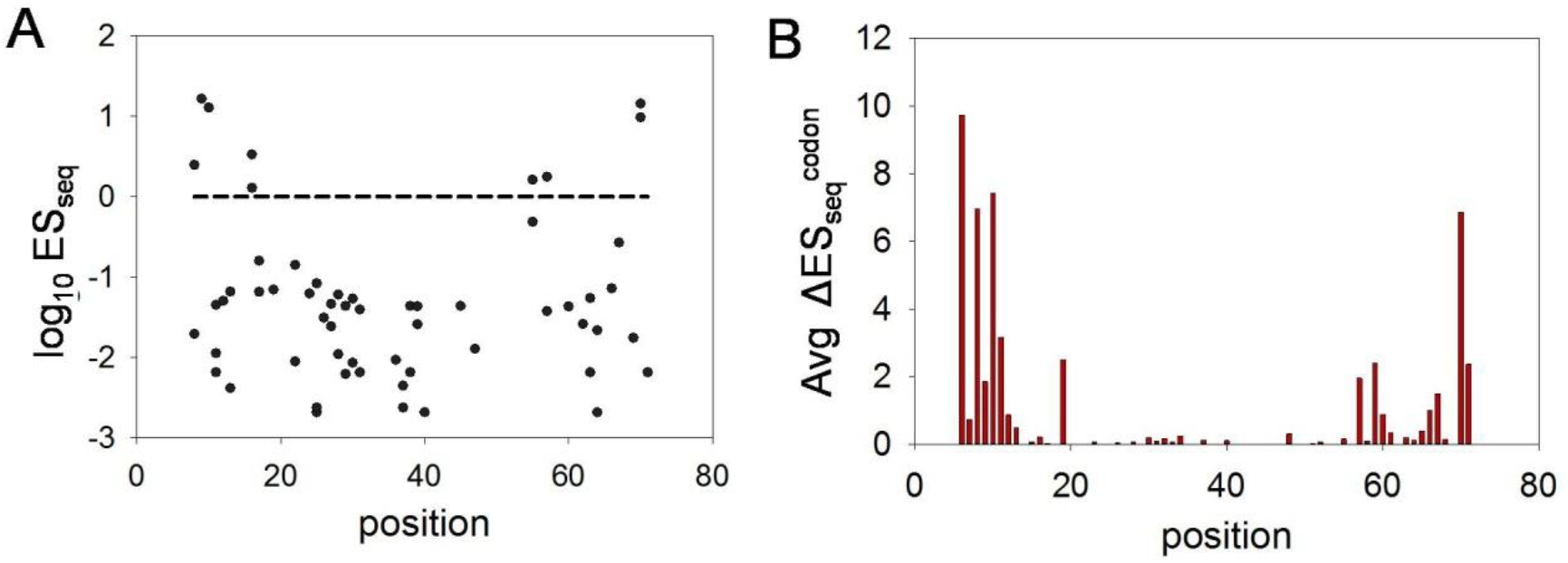
Inactive phenotype of synonymous mutants and codon specificity effects for non-synonymous mutants are non-uniformly distributed across the length of *ccdA*. (A) Distribution of log_10_ES_seq_ scores for all 62 available single synonymous mutants as a function of residue position. The WT ES_seq_ score of 1 (log_10_ES_seq_ = 0) is depicted as a black dashed line. Loss of function mutations with low ES_seq_ values are distributed across the length of the *ccdA* gene. A few mutants show WT like or higher ES_seq_ values and are largely clustered at the terminal regions of the gene. Phenotype of each synonymous mutant of CcdA is described in Table 2. (B) Averaged pairwise absolute differences in the ES_seq_ scores amongst codons as a function of residue position in CcdA. The differences between ES_seq_ values of each pair of synonymous codons encoding the same mutant amino acids are calculated. The average of the absolute differences of all mutant codon pairs for a residue position is estimated as follows: Average ΔES_seq_^codon^= (∑│ES_seq_^a^ - ES_seq_^b^│)/ ^n^C_2_, where a and b are different codons encoding a missense mutation and n is the total number of such codons at a position that have data for fitness scores. Regions close to the termini show the greatest variation in mutational effects, whereas the central region is largely populated by loss of function mutations.

**Table 2.** Mutational sensitivities of all available synonymous mutant codons at each residue position of CcdA library in operonic context. The codons (synonymous with respect to WT sequence of CcdA) have been colored based on the deep sequencing phenotypic activity (ES_seq_ scores) in operonic context. Dark green depicts hyperactivity (ES_seq_>1.5), while light green denotes WT like activity (0.7≤ES_seq_≤ 1.5). Yellow and red indicate slightly inactive mutants (0.1<ES_seq_< 0.7) and highly inactive mutants (ES_seq_≤ 0.1) respectively.

| Residue position | Synonymous mutant codons |  |  |
| --- | --- | --- | --- |
| 8 | ACG | ACT |  |
| 9 | GTG |  |  |
| 10 | GAT |  |  |
| 11 | AGT | TCG | TCT |
| 12 | GAT |  |  |
| 13 | AGT | TCG | TCT |
| 16 | CTG | CTT |  |
| 17 | CTG | CTT | TTG |
| 19 | GCT |  |  |
| 22 | GTG | GTT |  |
| 24 | ATT |  |  |
| 25 | AGT | TCG | TCT |
| 26 | GGG |  |  |
| 27 | CTT | TTG |  |
| 28 | GTG | GTT |  |
| 29 | AGT | TCG | TCT |
| 30 | ACG | ACT |  |
| 31 | ACG | ACT |  |
| 36 | GCG |  |  |
| 37 | AGG | CGG |  |
| 38 | AGG | CGG |  |
| 39 | CTT | TTG |  |
| 40 | AGG | CGG |  |
| 45 | AAG |  |  |
| 47 | GAG |  |  |
| 55 | GTG | GTT |  |
| 57 | AGG | CGT |  |
| 60 | GAG |  |  |
| 62 | AAT |  |  |
| 63 | GGG | GGT |  |
| 64 | AGT | TCG |  |
| 66 | GCG |  |  |
| 67 | GAT |  |  |
| 69 | AAT |  |  |
| 70 | CGG | CGT |  |
| 71 | GAT |  |  |

The mutational effects observed in case of synonymous mutations led us to investigate codon specific effects amongst non-synonymous CcdA mutants. We found several non-synonymous mutations at multiple positions having distinct phenotypes for different codons that code for the same mutant amino acid. Since NNK codons were used for mutagenesis and preparation of the SSM library, we could only investigate phenotypes of mutated codons having G/T at the 3^rd^ nucleotide position. Therefore, the phenotypic differences between codons could be investigated for a subset of mutated amino acid residues that are encoded by more than one codon, namely Alanine, Glycine, Leucine, Arginine, Proline, Threonine, Serine and Valine. We also found that phenotypic differences amongst codons in non-synonymous mutants were heightened in the N- and C-terminal positions of CcdA, while all mutations in the CcdA central stretch were inactive irrespective of the identity of the mutated codons as well as amino acids (Figure 2B). These results indicate the observed effects are not due to introduction of a mutated amino acid residue, but due to changes in the DNA sequences. The CcdA library under study comprises of a very small number of active mutants, which are both synonymous and non-synonymous and most are substantially more active in comparison to the WT construct (Figures 1C and 2). These active mutants were populated at the N and C-terminal residues, where mutations lead to the most diverse range of activities (Figure 2).

### Growth defects caused by *ccdA* mutations are not due to CcdB binding defects

The growth defects caused by *ccdA* mutations are only observed in the Top 10 strain that is sensitive to CcdB toxicity and not in the Top10 gyr strain (Supplementary Figure S4B), where the mutated Gyrase renders the strain resistant to CcdB mediated growth arrest. This clearly indicates that the growth defect displayed by *ccdA* mutations is caused by decreased inhibition of CcdB toxin in the sensitive strain. CcdA is known to rescue growth arrest by binding to CcdB with very high affinity, thus preventing CcdB from binding and poisoning DNA gyrase (Maki et al. 1996; De Jonge et al. 2009). Decreased inhibition could arise because of mutations affecting the folding of CcdA, mutations at residues in direct contact with CcdB, or because the total amount of expressed CcdA has been altered by mutation. In order to examine whether mutations significantly affect CcdA conformation, we characterized the ability of CcdA mutants to bind CcdB, using yeast surface display (YSD) methodology, where CcdA-CcdB binding can be decoupled from the toxic effect of CcdB on bacterial growth. CcdA was fused to *S. cerevisiae* Aga2p surface protein, displayed on the surface and its binding to purified CcdB toxin protein was measured using FACS (Figure 3). Although CcdA shows high mutational sensitivity throughout its length based on the phenotypic study in the operonic construct in *E. coli* (Figure 3A), when YSD CcdA libraries were constructed from plasmid DNA isolated from the sensitive and resistant *E. coli* strains by cloning in pETcon vector (Figure 3B), the YSD study revealed only minor differences in CcdB binding between these two libraries (Figure 3C). In addition, studies of individual mutants using YSD reveal that all CcdA synonymous mutants and WT protein bound equivalently to CcdB (Figure 3D), irrespective of the corresponding distinct mutant phenotypes in the native operonic context in *E. coli*. Synonymous mutations have previously been suggested to affect translation rate and co-translational folding, thus altering protein activity (Krasheninnikov et al. 1988; Komar et al. 1999; Sander et al. 2014). Such folding defects are unlikely in the case of CcdA, since its C-terminal domain, involved in CcdB binding is intrinsically disordered and remains natively unfolded in unbound conditions. A more extensive DMS study of CcdA mutants in the yeast surface display system indicates that mutations that affect the CcdB-binding affinity of CcdA primarily occur at residues directly involved in contact with CcdB (Chandra et al. 2022). While YSD fails to replicate the *in vivo* conditions of *E.coli* and coupling of transcription-translation to protein function, the binding assay on yeast surface clearly shows that most mutations do not alter the inherent CcdB-binding function of the intrinsically disordered CcdA molecule. Further, we find no discernable difference in the distribution of phenotypes or the ES_seq_ values between mutations at CcdB interacting and CcdB non-interacting residues in the C-terminal CcdB binding domain of CcdA, classified based on the available structure of CcdA C-terminal domain bound to dimeric CcdB (De Jonge et al. 2009) (Supplementary Figure S5A). These observations indicate that mutational effects on binding affinity to CcdB contribute insignificantly to the mutant phenotypes observed for *ccdA* in its native operonic context.

**Figure 3.**
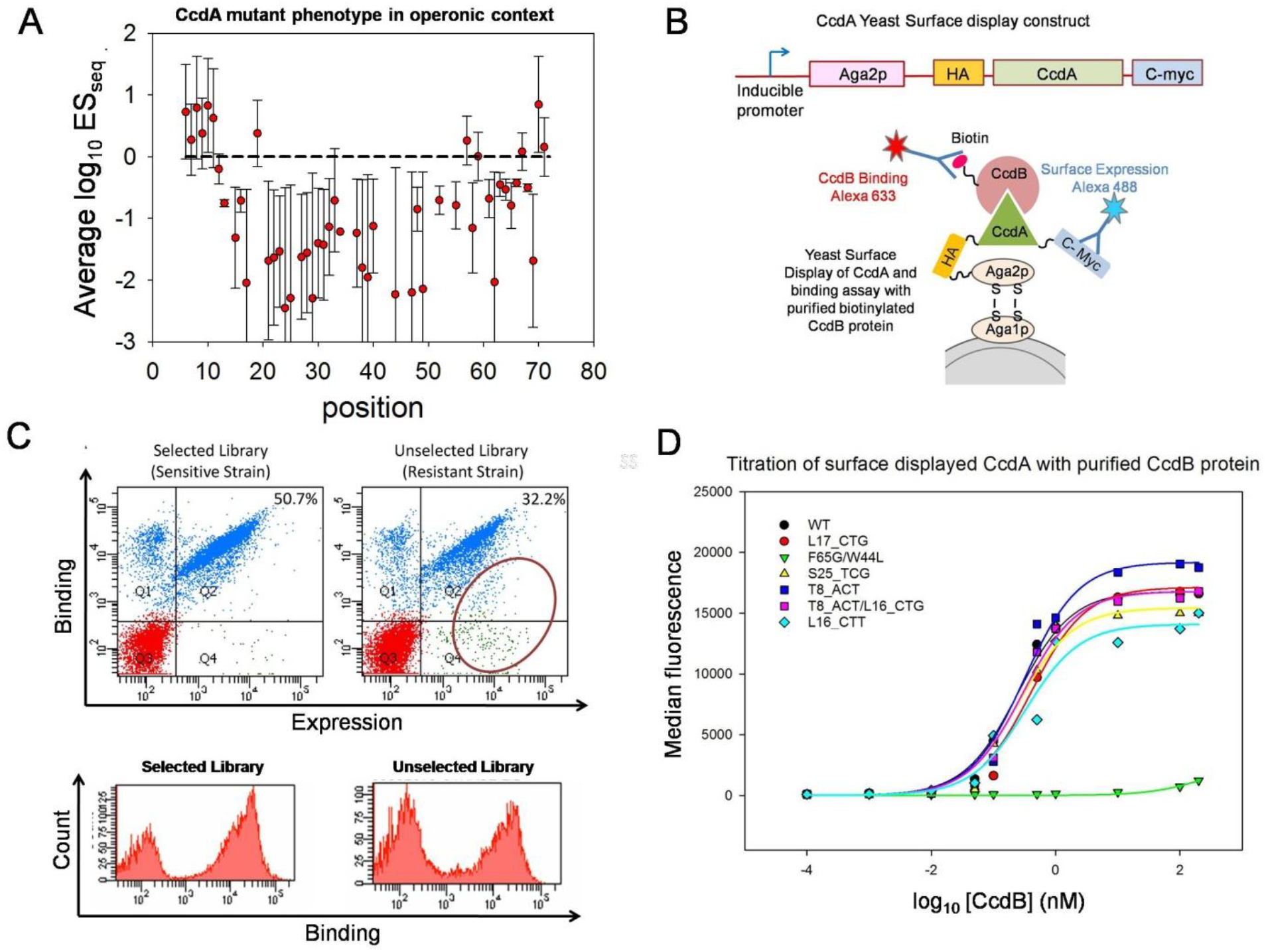
Observed growth defects in CcdA mutants are typically not caused by loss of CcdB binding activity. (A) Average log_10_ES_seq_ values (red circles) as a function of residue position. The error bars represent the associated standard deviation and the dashed line indicates the WT ES_seq_ value of 1. (B) Schematic representation showing (top) a monocistronic fusion construct of CcdA gene for (bottom) yeast surface display and subsequent probing of surface expression and CcdB binding using flow cytometry (*see Methods*). (C) The CcdA mutant libraries derived before and after selection in *E.coli* strains were cloned in the YSD vector, surface displayed and probed for surface expression and CcdB binding using FACS. Double plots and histograms are shown in the upper and lower panel respectively. The resistant strain derived library shows only marginally lower CcdB binding than the sensitive strain derived library, indicating that the number of CcdA mutants having significantly lower CcdB binding activity in comparison to WT CcdA is very low (population marked with red circle). Lower panel shows the CcdB-binding histograms of the two libraries studied using flow cytometry, which are not significantly different. (D) Binding of individual CcdA mutants to WT CcdB probed by yeast surface display. CcdB binding affinity was assessed by titrating the cells displaying an individual CcdA mutant with various concentrations of biotinylated CcdB, followed by binding to streptavidin conjugated AlexaFluor-633. Labelled cells were analyzed on a BD FACSARIA III. The binding affinity was calculated by fitting median fluorescence intensity to a one site binding model. All single-site synonymous mutants of CcdA (showing significant variation in phenotypes in the operonic study, see Table 1) showed comparable binding affinity to WT (see Table 3). The double mutant F65G/W44L was taken as a negative control that shows impaired CcdB binding.

**Table 3.**
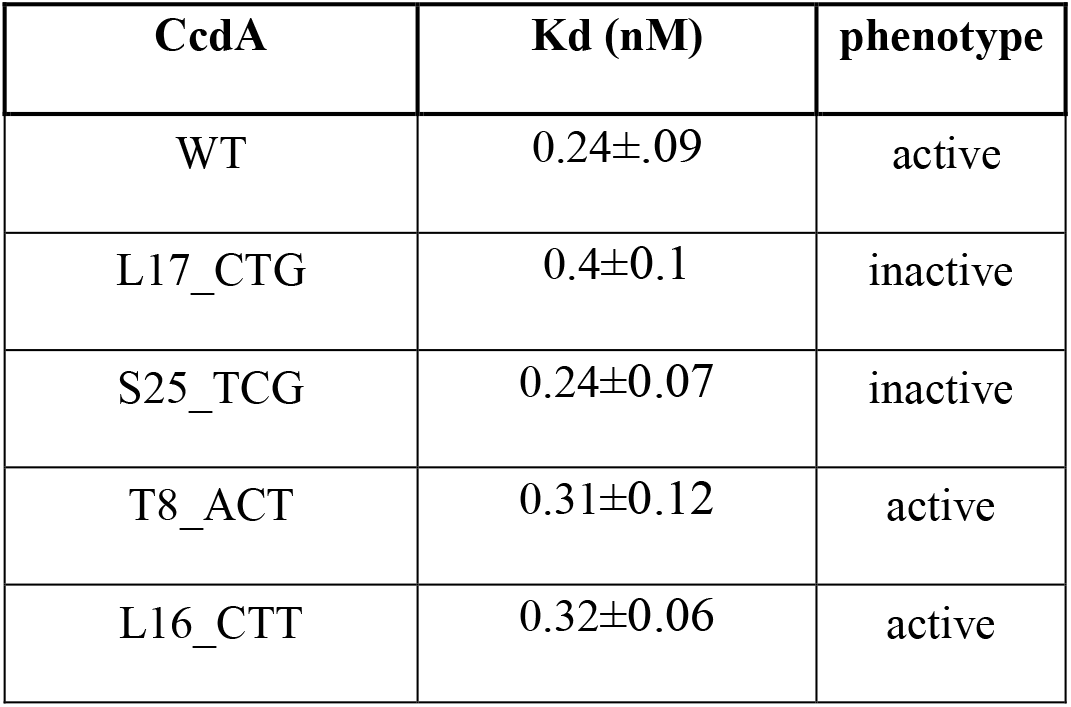
Binding affinities of CcdA mutants and WT calculated using titration of yeast surface displayed CcdA molecules with purified CcdB. The dissociation constants (K_d_) were obtained by fitting the fluorescence intensity titration traces to a single-site ligand binding model. In the operonic context in *E.coli,* the single codon synonymous mutants L17_CTG and S25_TCG mutants display an inactive phenotype, while T8_ACT and L16_CTT have an active phenotype based on the deep sequencing studies.

We also studied the distribution of mutational phenotypes (for non-synonymous mutations) at different classes of functionally important residues in the N-terminal domain of CcdA, identified based on the available NMR structure of the DNA bound dimeric CcdA N terminal domain (Madl et al. 2006). We found that mutations in the DNA binding residues in the CcdA N-terminal domain have no distinguishable difference in terms of the growth phenotype, when compared to other non-interacting exposed residues (Supplementary Figure S5B). Surprisingly, mutations at buried positions in the N terminal domain show significantly reduced activity and more severe growth defects, especially in the case of mutations to non-aliphatic residues, when compared to any other classes of CcdA residues (Supplementary Figure S5C). This suggests that mutations that can disrupt the N-terminal buried core of CcdA can induce increased protein degradation or proteolysis, ultimately producing the observed inactive phenotypes.

### *E. coli* genomic codon usage contributes to growth phenotypes of single codon mutants in CcdA

The frequency of different degenerate codons (for any amino acid) in the genome of an organism varies significantly. We investigated the relationship between the phenotypes of mutants and relative codon usage (in the *E.coli* genome) of the introduced codon. The distribution of relative codon usage (RCU) values amongst active and inactive CcdA mutants was significantly different (Figure 4A). It has been suggested that having a higher percentage of frequent codons in a gene leads to higher amounts of expressed protein, across various kingdoms of life (Sharp et al. 1986; Shields and Sharp 1987; Sharp and Devine 1989; Karlin et al. 1998). Decreased levels of CcdA protein *in vivo* will result in the presence of unbound CcdB molecules that will poison bacterial DNA Gyrase, resulting in cell death. While there is a large fraction of inactive mutants in the *ccdA* single mutant library, we observe that ∼70% of *ccdA* mutants displaying an active phenotype harbor frequent codons (based on *E.coli* genome) with similar or higher codon usage frequency relative to the WT codon. The observed high mutational sensitivity in the central region of the *ccdA* gene might result from the high prevalence of frequent codons in this region (Supplementary Figure S6A). Thus, most mutations at these positions lead to the introduction of rarer codons, possibly leading to lower CcdA protein expression and subsequent cell death. However, the distribution of relative tRNA abundance values amongst active and inactive mutants was similar (Figure 4B). There was no statistically significant difference between inactive and active mutant distributions for other sequence features such as relative GC content and number(s) of base changes upon mutation. (Figure 4C-D).

**Figure 4.**
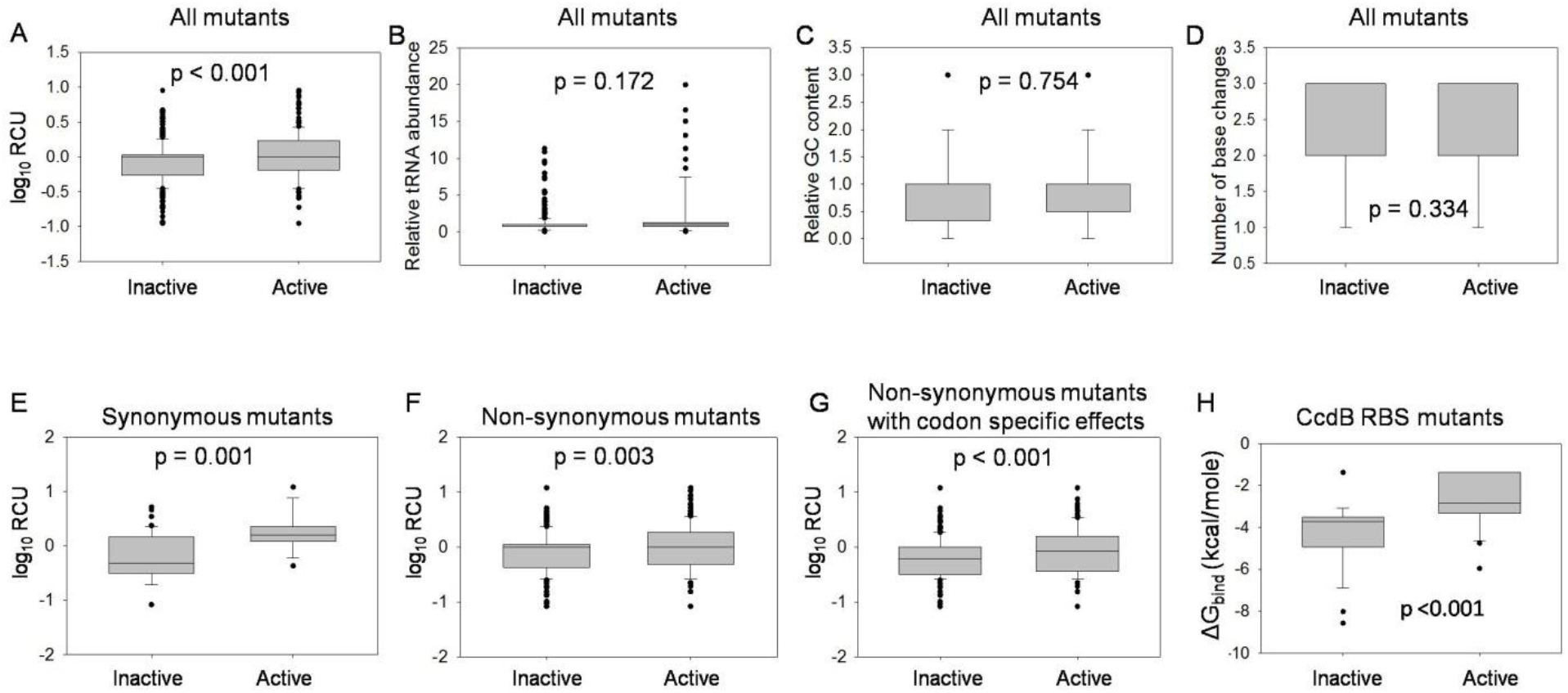
Relation between DNA sequence features and growth phenotypes of CcdA mutants. All available mutants in the CcdA library (N=1528) were divided into active (N=314) and inactive (N=1214) classes based on an ES_seq_ score cut-off of 0.1. Mutant distribution was plotted with respect to various sequence based features, namely relative codon usage RCU (A), relative tRNA abundance (B), relative GC content (C) and number of base changes upon mutation (D) calculated for mutant codon with respect to WT codon (*See Methods*). The distribution of sequence feature scores for inactive and active mutants have been plotted as box plots, where the central straight line depicts the median, the box the interquartile range (IQR), edges of the box the first and third quartile (Q1 and Q3), the whiskers minimum (Q1-1.5IQR) and maximum (Q3+1.5IQR) of the distribution and the black circles the outliers. Statistical significance was assessed by a Mann-Whitney Rank Sum Test. A p-value >0.05 indicates that there is no statistically significant difference between the distributions of the indicated parameter for active and inactive mutants, at a 95% confidence interval. Only the distribution of RCU is significantly different across the inactive and active class of mutants. The mutant distribution with respect to RCU was also plotted for *ccdA* synonymous mutants (N=62) (E), non-synonymous mutants (N= 1466) (F) as well as the mutants showing codon specific effects (N=550) (G), all of which showed significant differences in distributions of active and inactive classes. CcdA non-synonymous mutants encoded by different codons (for mutated amino acids encoded by more than one codon) showing different phenotypes for the different codons are referred to as mutants showing codon specific effects. (H) Effects of mutations in the CcdB RBS site in *ccdA* gene on phenotype in the native operonic context. The free energy of binding to the anti-Shine Dalgarno sequence on the ribosome, denoted as ΔG_bind_ (CcdB RBS) was calculated for each of the possible single codon mutations at the CcdA residue positions 70 and 71 that constitute the ribosome binding site (RBS) of the downstream CcdB gene. Calculations were done using the RNAsubopt program in the Vienna RNA package. Distribution of ΔG_bind_ (CcdB RBS) values for inactive and active mutants at residue positions 70 and 71 of CcdA are significantly different. Statistical significance was assessed by a Mann-Whitney Rank Sum Test.

We also studied the distribution of these DNA sequence features among the active and inactive variants for the subsets of synonymous mutations, non-synonymous, as well as for the non-synonymous substitutions which showed codon specific effects. Inactive mutations show significantly lower relative codon usage values than active mutants in all subsets consistently (Figure 4E-G).

Codon Adaptation Index (CAI) is a commonly used parameter that describes the degree of codon bias for a whole gene or sequence and helps to infer the relative adaptiveness of the sequence (Sharp and Li 1987). While it is routinely used in studies that investigate functional changes in sequences where multiple rarer or more frequent codons have been introduced, a single-site mutation (change of one codon) does not produce a large change in the CAI. CcdA WT gene has a CAI of 0.76 and CAI calculated for single codon changes are in the range of 0.7-0.78. Such small changes are not expected to show any distinguishable effects on relative fitness of mutants. We find that distribution of relative CAI values (mutant CAI/WT CAI) across inactive and active mutants fail to show any significant difference in the Mann Whitney test (Supplementary Figure S7). On the other hand, relative codon usage (RCU) uses the relative fraction of usage of the synonymous codon encoding a particular amino acid and also takes into account the most common codon (for the WT amino acid). Estimating phenotypic effects due to altered synonymous codon usage fraction at the particular site using RCU helps to reveal that codon usage bias does indeed play a role in affecting the fitness in single-site mutants as described earlier (Figure 4A, D-F).

### Strength of the CcdB ribosome binding site determines with CcdA mutant phenotype

A primary sequence element that is known to be a strong determinant of the efficiency of translational coupling is the Shine Dalgarno (SD) sequence. Early studies by Das and Yanofsky in the Trp operon showed that translation initiation at any start site located near a functional stop codon of the upstream gene may be influenced by the strength, sequence and location of the SD region and its spacing from the start codon (Das and Yanofsky 1989). The Ribosome Binding Site (RBS) for the *ccdB* gene lies within the 72 amino acid CcdA coding region (*ccdA* gene), primarily spanning residues 70-71. Therefore, mutations in these residues may affect the relative CcdA and CcdB levels by changing the strength of the RBS for CcdB. An analysis of all available codon substitutions at residues 70 and 71 of CcdA from the deep sequencing data indicated that the predicted strength of the RBS for CcdB expression was a significant contributor to the phenotype of these mutants (Figure 4H). Mutants at residues 70 and 71 displaying inactive phenotype were predicted to improve the CcdB RBS strength. The molecular basis of one such mutation, D71R was examined. Since two of the arginine codon mutants had an active phenotype, it is unlikely that the inactive phenotype observed for the AGG arginine codon is due to effects on protein stability or protein-protein interaction. It is known that the consensus sequence for the RBS, having the maximal strength is 5’ AGGAGG 3’ (based on the corresponding complementary sequence of 16S rRNA, 5’CCUCCU 3’). Therefore, it is likely that a change from the suboptimal RBS for CcdB present in the *ccdAB* operon (5’AGGGAC 3’), to a consensus RBS (as is predicted in case of D71R_AGG mutation), decreases the CcdA: CcdB ratio in the cell, there by leading to cell death. This prediction was consistent with data from quantitative proteomics (Figure 5A).

**Figure 5.**
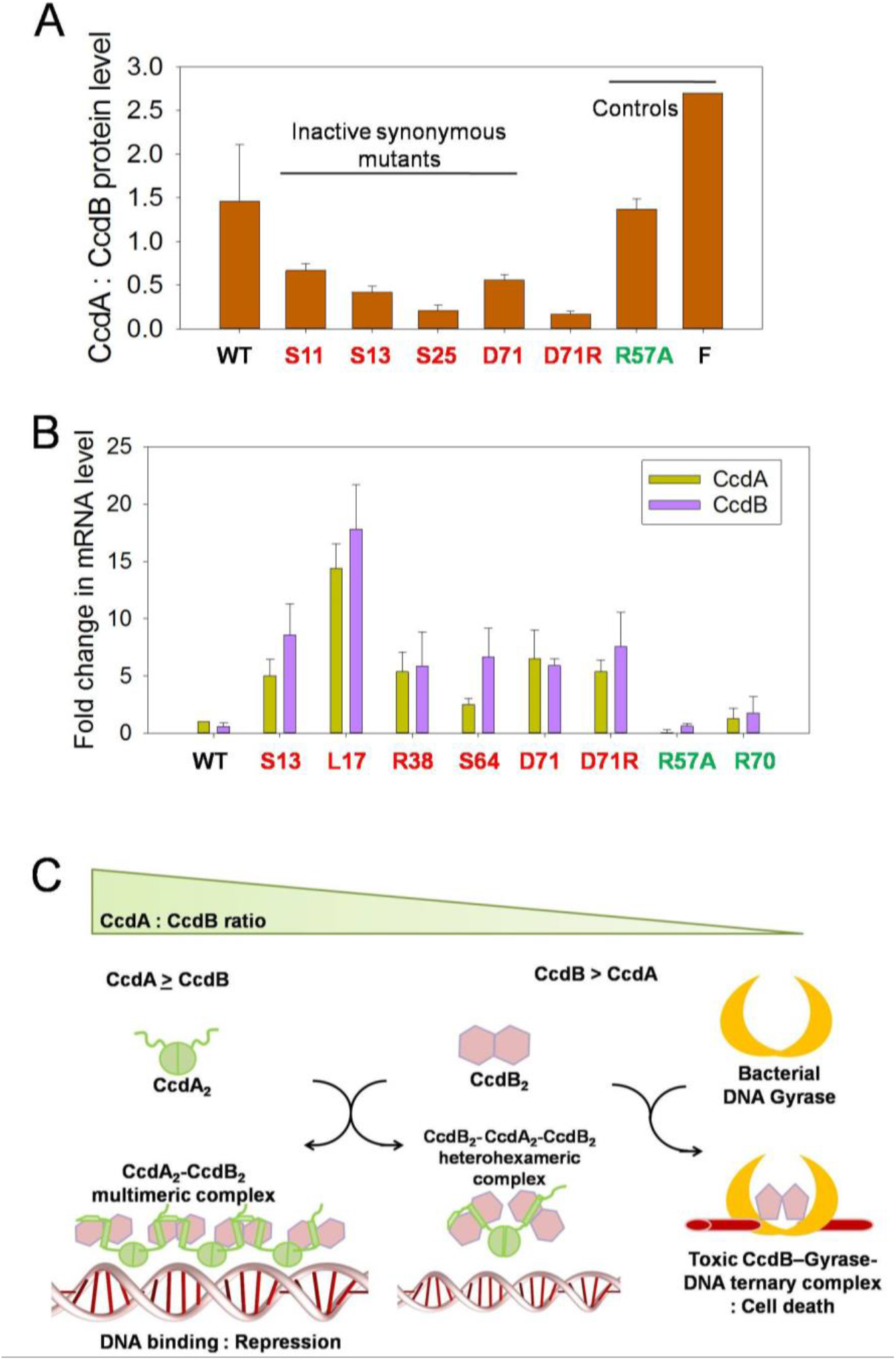
Decreased CcdA: CcdB protein ratio leads to loss of function phenotype for inactive synonymous CcdA mutants and also increases *ccdAB* mRNA levels by feedback autoregulation of the operon. (A) Relative levels of CcdA and CcdB peptides in *E. coli* (Top 10 gyr strain) lysates were determined for WT as well as selected single synonymous mutants of CcdA using a quantitative proteomics approach. All synonymous inactive mutants studied have a decreased CcdA:CcdB protein ratio. The inactive non-synonymous mutant D71R which is predicted to improve CcdB translation due to a stronger CcdB RBS, also displays a decreased CcdA: CcdB protein ratio relative to WT. (See Table1 for phenotypic scores of mutants). F indicates the value for F-plasmid. Mutants showing active and inactive phenotypes in the operonic study are marked in green and red respectively. (B) Inactive synonymous mutants show higher levels of *ccdAB* mRNA, while active mutants have lower or WT comparable levels of *ccdAB* mRNA in Top10 gyr *E. coli* cells. The mRNA levels of *ccdAB* specific transcripts were determined using q-RTPCR. *ccdA* and *ccdB* regions were both independently quantitated. Mean Ct represents the threshold cycle for amplification obtained from duplicate samples from two different experiments. Fold change in *ccdA* and *ccdB* transcript levels are with respect to the WT Ct values (see Methods). Error bars indicate estimated standard deviation of the measurements. The mutants are labeled as WT amino acid identity_residue position_mutated amino acid identity in case of non-synonymous mutants and as WT amino acid identity_residue position in case of synonymous mutants. (C) Schematic model for CcdA:CcdB ratio dependent autoregulation of *ccdAB* operon transcription as described previously (Vandervelde et al. 2017).

### Mutations in CcdA lead to altered levels of CcdA proteins and therefore modified CcdA:CcdB ratio in cell

Codons for inactive CcdA mutants have significantly lower relative codon usage in comparison to active mutants, suggesting that the observed mutational phenotypes might result from alterations in translational rate of the CcdA or CcdB gene products. To further probe protein levels in mutants, we carried out a proteomics study. The extremely low levels of CcdA and CcdB proteins *in vivo* cannot be accurately characterized using classical methods like SDS-PAGE or Western Blotting. The ratio of CcdA: CcdB in WT was found to be slightly larger than 1 in the proteomics study (Figure 5A). Interestingly, all the five inactive CcdA mutants tested show a lower CcdA:CcdB ratio (Figure 5A). Introduction of rarer codons thus appears to reduce the translation of CcdA, decreasing the CcdA:CcdB ratio of proteins in the cell, causing more severe growth arrest. The inactive D71R_AGG mutant (CcdB RBS site mutation in *ccdA*) also exhibits a lower CcdA:CcdB ratio, likely due to increased translation of CcdB protein. Active mutant R57A and CcdAB operon in F-plasmid show comparable protein ratio relative to WT.

### Increased levels of operonic mRNA in inactive mutants confirm decrease in CcdA:CcdB ratio

The five inactive and one hyperactive mutants of CcdA tested in the proteomic study have relatively reduced and elevated levels of CcdA proteins relative to WT respectively. The *ccdAB* expression at the transcription level is autoregulated by the CcdA-CcdB protein complex in a CcdA:CcdB ratio dependent manner (Tam and Kline 1989; Afif et al. 2001). When the CcdA:CcdB protein level ≥l, elongated multimeric CcdA-CcdB complexes with alternating dimeric CcdA_2_ and CcdB_2_ are formed which bind to the *ccdAB* operator region with high affinity, repressing gene expression. On the other hand, when CcdA:CcdB <1 (protein level), hetero-hexameric complexes of one CcdA dimer bound to two CcdB dimers (CcdB_2_-CcdA_2_-CcdB_2_) are formed that have low binding affinity to the operator region and leads to a derepressed state of operon, allowing transcription to proceed (Vandervelde et al. 2017).

To study the possible effects of CcdA sequence changes resulting from single codon mutations on gene expression, we quantified *ccdAB* operon specific RNA using RT-qPCR using both *ccdA* and *ccdB* gene specific primers. Relative amounts of *ccdA* and *ccdB* encoding transcripts were near identical in WT and were not significantly changed in case of most mutations (ratio of *ccdA* and *ccdB* specific mRNA levels ≈ 1) (Figure 5B), as expected for transcription from an operon. However, significant increases in the mRNA levels for both the *ccdA* and *ccdB* amplicons were observed in case of all inactive mutants (Figure 5B). This is consistent with the lowered CcdA:CcdB protein levels in the inactive mutants observed in the previous section. A lowered CcdA:CcdB protein ratio will result in de-repression of the operon, leading to upregulation in mRNA production (Figure 5C). Analysis of whole genome RNA-seq and ribosome profiling data available for the *E.coli* genome indeed indicate that while the mRNA levels for most type II TA systems were similar for the toxin and the antitoxin consistent with our observations in *ccdAB*, the antitoxin protein levels can be upto two fold higher than the toxin under normal growth conditions(Deter et al. 2017). This indicates that differential translation of the *ccdA* and *ccdB* genes and/or differential proteolysis of the two proteins play a vital role in maintaining the toxin-antitoxin protein levels *in vivo*. Altered DNA sequences affecting the relative levels of the toxin-antitoxin proteins can thus be detrimental to functions of TA systems.

### Investigation of possible effects of ribosomal pausing on CcdA mutant phenotype

An analysis of bacterial genomes has suggested that Shine Dalgarno (SD) like sequences within the coding region of genes could be potential ribosomal pause sites and are therefore under negative selection (Li et al. 2012; Mohammad et al. 2016). To examine if mutations in CcdA generate such potential pause sites, the difference in interaction energies with the *E. coli* ribosomal anti-SD sequence between single synonymous mutants and the WT sequence was calculated for a window of 10 nucleotides using the RNAsubopt program in the Vienna RNA package (Gruber et al. 2008). The data indicate that the inactive and active synonymous mutations show no distinguishable difference in anti-SD interaction (Supplementary Figure S8A). The extent of ribosome pausing at each codon and the associated effects on *in vivo* translation speed have been estimated previously (Chevance et al. 2014). In the present study, we did not observe any significant differences in distribution of these relative ribosome stalling values for different codons between synonymous inactive and active mutants of CcdA (Supplementary Figure S8B).

### Effect of single codon synonymous mutations in another TA module, RelBE

To test if other TA modules are also sensitive to synonymous mutations, the RelBE system was used as a test case. Unlike the plasmidic *ccdAB* system, *relBE* is natively a chromosomal toxin-antitoxin system, expressed as a part of the *relBEF* operon (Cherny et al. 2007). The RelE toxin is a site-specific, ribosome dependent mRNA endonuclease. Expression of free RelE leads to bacterial growth inhibition. Mild overexpression of RelE is also reported to increase persister cell frequency (Fasani and Savageau 2013). While unrelated in sequence, the RelB protein from *E. coli* is similar to the CcdA antitoxin in both fold and thermodynamic properties, and inhibits the action of RelE by forming a tight complex. The RelBE complex acts as a transcriptional repressor and autoregulates its own expression (Cherny et al. 2007). Homologs of this system are found in many bacterial species including pathogens like *Haemophilus influenzae.* An analysis of ribosome profiling data for the *E. coli* K12 MG1655 strain indicates that the translation efficiency for the *relB* gene is 3.6-fold higher than the *relE* gene (Deter et al. 2017), although both the codon and tRNA adaptation indices are similar for both genes, similar to what we observed in the CcdAB system.

To study mutational effects, *relBE* operon was cloned in pUC57 plasmid. We studied five single codon synonymous mutations, distributed across the length of the *relB* gene in a native *relBE* operonic context, including both three and two base substitutions. These mutations were to both more frequent and rarer codons. A control mutation at residue 77 of *relB* that strengthens the SD sequence of RelE was also studied. This is expected to lead to an increase in RelE levels and inhibition of cell growth (Figure 6). Using a similar screen of cell death versus growth as in the case of CcdAB system, we monitored the growth of single synonymous mutants in the WT (*E. coli* BW25113) as well as in the toxin deleted (*E. coli* BW25113 Δ*relE*) strain. The two *relB* variants with rarer mutated codons tested here had a decreased growth in the WT strain (Figure 6B-C). We also found that two mutations of *relB* with higher codon usage with respect to WT codon display improved growth phenotype. The growth defective D77 variant involves introduction of a rarer codon besides strengthening the RBS. The *relB* mutant phenotypes indicate the existence of a relatively general phenomenon wherein a change in DNA sequence upon synonymous mutations can have observable phenotypic effects for toxin-antitoxin operon systems, where small perturbations in the system can compromise cellular fitness.

**Figure 6.**
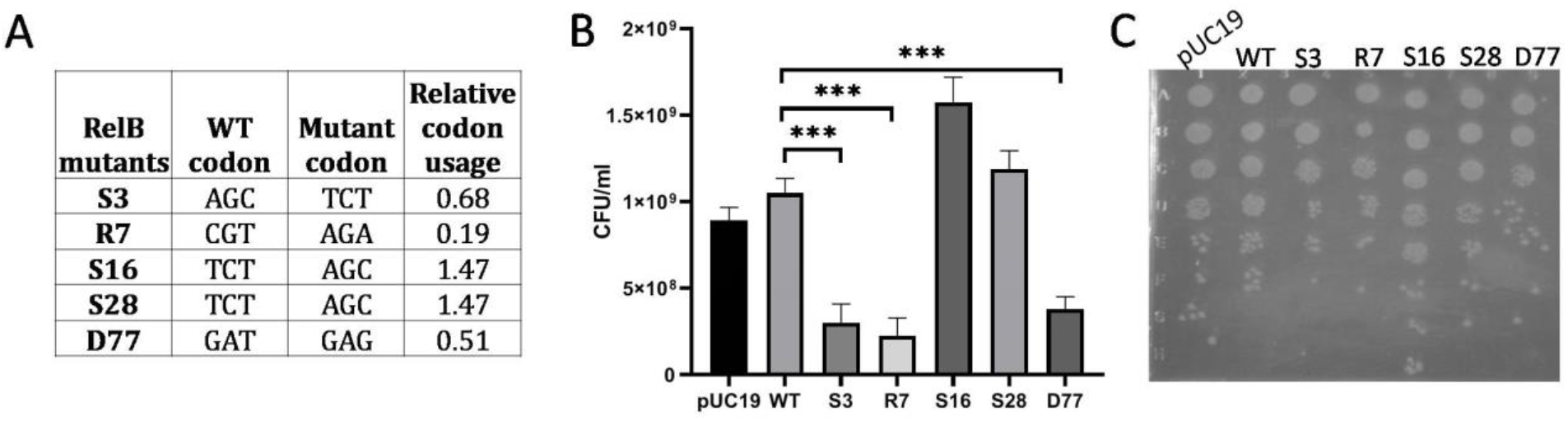
Phenotypic effects of select single synonymous substitutions in *relB* antitoxin gene of the RelBE operon. (A) List of various synonymous mutations made in the *relB* gene in the native operon context along with their relative codon usage upon mutation with respect to the WT codons. A single base substitution at the D77 residue of RelB which increases the strength of the SD sequence for downstream toxin RelE was used as a control mutant. D77 mutant also involves introduction of a rarer codon. (B-C) Growth of single synonymous mutants in *E. coli* BW25113, sensitive strain. Following transformation, ten-fold dilutions were plated (B) as well as spotted (C) on LB-Agar plates. The number of colonies formed in the dilution plate with ∼50-100 colonies were counted for each mutant and CFU/ml was calculated and reported. (B) The D77 synonymous mutant, expected to increase the levels of the downstream RelE toxin, displays an inactive phenotype. RelB synonymous mutants with rarer codons display decreased growth, while frequent codon bearing mutants exhibit higher growth relative to WT. The error bars indicate standard error. ‘*’ indicate significant differences from the WT (*** p value <0.001, paired t-test).

## Discussion

In type II TA systems, the toxin and antitoxin can form higher order TA complexes which bind to the TA promoter repressing its transcription. Conditional co-operativity (T:A protein ratio dependent co-operative interaction of TA complex with the operator DNA) is known to be a dominant mechanism in regulating the transcription of most TA systems (Cataudella et al. 2012). Given the higher rate of degradation for the antitoxin due to its disordered nature, the antitoxin needs to be produced at a higher rate than the toxin, for the cells to be viable. However, mechanisms that can lead to differential levels of the toxin and the antitoxin protein from a common mRNA are not well understood. While protein levels can in principle be regulated at both the transcriptional as well as the translational level, RNA-seq data from *E.coli* indicate that mRNA levels for the toxin and the antitoxin are not significantly different, as is expected for genes encoded by a single polycistronic mRNA (Deter et al. 2017). Ribo-seq data available for TA systems indicate that the protein synthesis rates (calculated from ribosome densities) for the antitoxin are at least two fold higher than the toxin, for six TA pairs for which sufficient data is available (Deter et al. 2017). These observations indicate that there are indeed nucleotide sequence dependent features, which allow regulation at the level of translation, leading to differential synthesis rates for the two genes in the operon. Maintaining a perfect balance between the antitoxin and toxin proteins is essential for proper functioning of TA modules. Small changes in gene expression caused by altered DNA sequences, both in coding or non-coding regions can potentially be detrimental to this balance and result in observable phenotypes. In the present study, we examined mutational effects in the bacterial *ccdA* antitoxin gene on phenotype, in its TA operonic context, using saturation mutagenesis coupled to deep sequencing. We observe significantly high fractions (∼80%) of both synonymous and non-synonymous mutants to show reduced antitoxic functions in the operonic context, leading to reduced survival of the bacterial host cells. An extensive review by Bailey et al (Bailey et al. 2021) summarizes the DFE’s of synonymous mutations for twelve different proteins and viral genomes. Studies investigating distribution of fitness effects in bacterial and yeast proteins have revealed that most mutants (>95%) retain at least 50% fitness (growth-rate based) with respect to WT (Lind et al. 2010; Schenk et al. 2012; Lebeuf-Taylor et al. 2019). Similar results are seen in a comprehensive study of Hsp90 (Flynn et al. 2020). Other studies revealed DFE of missense mutants to be bimodal. For bacterial TEM-1 (Firnberg et al. 2016), ∼25% missense mutations showed loss-of-function and another ∼30% showed reduced fitness. For yeast Hsp90, ∼50% missense mutations showed 10-50% reduced fitness (Hietpas et al. 2011). In case of various viral genes/genomes 20-40% mutants show lethal (loss-of-function) and 10-35% show deleterious (reduced fitness) phenotypes (Carrasco et al. 2007; Domingo-Calap et al. 2009; Peris et al. 2010; Wu et al. 2014). Notwithstanding these variations in DFE for missense mutants, all previous studies indicate most synonymous mutants to be near neutral with the exception of cases where only a small number of synonymous mutants displays a range of reduced (20-70%) fitness (Carrasco et al. 2007; Lind et al. 2010; Cuevas et al. 2012; Wu et al. 2014). For *ccdA*, we observe that 80% of both synonymous and non-synonymous *ccdA* mutants are observed to have ES_seq_ value ≤ 0.1 (ten fold lower colonies obtained following transformation into Top10 *E.coli* strain relative to WT) in the present study. The major peak in the distribution of ES_seq_ scores in *ccdA* mutants is shifted towards lower values relative to the WT. While ES_seq_ is not identical to conventional fitness measures, it is still apparent that the fraction of synonymous (as well as non-sysnonymous) mutations showing significantly reduced activity is higher than those seen in the other systems, some of which are summarized by Bailey et al. Most importantly, we observed that most inactive *ccdA* mutants displaying ES_seq_ < 0.1 produced 10-1000 fold lower number of colonies than WT *ccdA* when transformed in the CcdB-sensitive *E.coli* strain Top10 (but similar numbers of colonies when transformed in toxin resistant strain Top10 gyr to WT). In the previous DFE studies involving bacterial/yeast growth-rate based phenotypic investigation, only mutants with comparable number of colonies to ancestor/WT sequence retrieved after transformation were used for further DFE investigations (Lind et al. 2010; Bailey et al. 2014; Lebeuf-Taylor et al. 2019). This indicates that the *ccdAB* operonic system under study is apparently more sensitive to mutations than those previously studied. We also found the *ccdAB* operonic system under study was more sensitive to non-synonymous mutations than synonymous mutations in the *ccdA* gene. A reduced severity in fitness defects in case of synonymous relative to non-synonymous mutations is also observed in few other cases (Fragata et al. 2018; Bailey et al. 2021), though the overall mutational sensitivity in these studies was found be much lower than observed for CcdA.

The *ccd* operon used in the present experiments differs slightly in sequence from the one on F-plasmid (Supplementary Figure S1), resulting in a slightly reduced CcdA: CcdB protein ratio (Figure 5A). This can be attributed to the presence of a suboptimal *ccdA* RBS in the non-coding region (Supplementary Figure S1) in our WT CcdAB construct. Small decreases in the ratio caused by mutations have dramatic phenotypic effects. This results from the very high toxicity of CcdB, the feedback regulation present in the system and the additional amplification resulting from use of a high copy number pUC plasmid to house the *ccd* operon. The readout here is a function of the CcdA:CcdB ratio. While it is very challenging to measure absolute levels of either protein, we are able to measure the ratio using proteomics. The data in Figure 5A show, consistent with the observed phenotypes, that the CcdA:CcdB ratio is <1 for the four inactive synonymous mutants tested, whereas it is ∼1.4 in the WT operon. For reasons that are currently unclear, in the *ccd* operonic construct we have used, mutations that enhance the CcdA:CcdB ratio and result in higher values of ES_seq_ (indicating beneficial variants) are rare, in contrast to other systems described in Bailey et al (Bailey et al. 2021) and Flynn et al (Flynn et al. 2020). The present *ccd* construct amplifies effects of point mutations and allows detection of changes that would be missed in other systems where small changes in expression level do not result in observable phenotypes. Strong mutant phenotypes were also found for single-site synonymous variants of the *relB* gene in RelBE TA operon, also cloned in a high copy number plasmid. Single codon changes in *relB* produced observable changes in phenotype comparable to those in the *ccdAB* system. The current study therefore showcases the utility of TA operons for providing general insights into the plethora of mechanisms by which point mutations can affect phenotype.

The present studies reveal that most single codon substitutions, including synonymous mutations along the length of *ccdA* lead to a loss of function phenotype. Many non-synonymous mutations also show codon specific phenotypes, meaning that significantly different phenotypes are observed upon introduction of different codons that encode the same mutated amino acid residue. These studies demonstrate that there can be strong selection for specific synonymous codons. This complicates interpretation and use of dN/dS ratios for studying selection pressure especially in operons. The observed dependence of the mutant phenotype on relative codon usage of mutant with respect to WT suggests that single codon mutations in CcdA likely influence the translation efficiency of the CcdA, thus altering the relative levels of CcdA and CcdB proteins in the cell. A high frequency of frequent codons in the central region of *ccdA* gene suggests that the observed high mutational sensitivity at this region is because any codon introduced upon mutation is expected to be rarer relative to the WT codon. At the *ccdA* gene extremities there is larger variation in codon usages values (Supplementary Figure S6). Hence mutated codons at these regions may be more or less frequent relative to WT. Altered DNA sequence can directly affect ribosome assembly, ribosome stalling, rate of translation, or promote altered mRNA structure, which in turn can have multifaceted consequences on protein production. The quantitative proteomics study reveals that for loss of function mutants, the *in vivo* CcdA: CcdB protein level ratio is lower than that of WT, resulting in cell death. Codon usage has previously been found to be a determinant of protein translation efficiency, accuracy as well as kinetics (Ikemura 1985; Sørensen et al. 1989; Quax et al. 2015; Lyu et al. 2020). Most prior studies do not characterize the change in protein levels associated with single synonymous codon substitutions. However, total functional protein levels have been found to increase in case of beneficial single-codon synonymous mutants in TEM-1 by upto 6-7 fold (Zwart et al. 2018) and in ADAMTS13 enzyme by 1.4 fold (Hunt et al. 2019), without detected effects on transcription levels or protein conformation. An earlier study of an operonic system has revealed that synonymous mutations alter relative protein levels of operonic gene products upto 3 fold, causing altered growth rates (Kristofich et al. 2018). *In vivo* translation rates of glutamic acid codons (GAA and GAG) were measured to be significantly different (and correlated to respective codon usage), despite both codons being read by same tRNA (Sørensen and Pedersen 1991). In FAE and CAT enzymes, protein levels were found to be reduced ina number of variants containing multiple synonymous substitutions across the genes (Amorós-Moya et al. 2010; Agashe et al. 2013). A study of introduction of a small number of rarer codons in the most highly expressed genes in *E. coli* also suggested that codon usage of a gene can affect the translation rate of gene with altered codons as well as of other genes due to perturbation of cellular tRNA supply (Frumkin et al. 2018).

The prevalence of large phenotypic variations amongst CcdA mutants exclusively at the N-terminal (first 20 residues) and C-terminal residues (56-72) might also be a result of mutational effects on translation initiation of *ccdA* and *ccdB* genes respectively. Mutations near the *ccdA* gene terminus, at residue positions 70 and 71 (that also constitute the ribosome binding site of CcdB) appear to affect translational efficiency of the downstream CcdB gene. Mutants at these positions which are predicted to show improved ribosome binding exhibit a lower *in vivo* CcdA:CcdB protein ratio and higher toxicity in *E. coli*. Mutations that decrease the CcdA:CcdB ratio result in de-repression of the operon, this is confirmed by the observed increased *ccd* mRNA expression levels in these mutants. The elevated operonic mRNA level is expected to amplify the mutational effect on differential translation efficiency of CcdA and CcdB.

Earlier studies which observe a strong influence of codon rarity and codon pair bias on protein level (Quax et al. 2015; Boël et al. 2016) typically used over-expressed proteins. In natural contexts, many bacterial genes are organized in operons to ensure co-regulation of genes involved in similar pathways and functions. In the special case of toxin-antitoxin operons, fine tuning of the relative amounts of antitoxin andtoxin is essential and directly linked to bacterial fitness. Given the abundance and diversity of these TA systems in many bacterial species, including the pathogen *Mycobacterium tuberculosis*, it is reasonable to assume that the cell must have evolved multiple mechanisms to ensure their stringent spatial and temporal regulation. We find that these relative protein levels can be perturbed through changes in DNA sequences. Although effects of synonymous mutations are often attributed to co-translational folding defects and altered activity of protein, we find that most of the inactive *ccdA* mutants are not CcdB binding defective, based on yeast surface display studies. We also observed that single mutations in DNA interacting residues of the CcdA N-terminal domain did not affect growth phenotype more drastically than mutations in other residues in CcdA, indicating no significant loss of DNA binding in such mutants. In contrast however, the heightened inactivity was observed in case of mutations at buried positions in the structured N-terminal domain. This suggests probable disruption of the hydrophobic core in the N-terminal domain, leading to increased proteolysis and decreased CcdA protein level. Besides the amino acid substitution related effects observed in the N-terminal domain, we also observe large phenotypic variation in the *ccdA* activity amongst a subset of missense mutants coded by different codons in both the N and C terminal regions of CcdA that cannot be explained by the codon usage bias or tRNA abundance. Interestingly, these effects occur in spans of about 40 bases which is on the order of the footprint of a single ribosome on mRNA in prokaryotes(Mohammad et al. 2019).Thus, it is possible that the high diversity in phenotype for synonymous codons exclusively at the two termini of *ccdA*may be caused by altered translation initiation rates for CcdA and CcdB in the operon with point mutations at the N- and C-terminus positions of the *ccdA* gene respectively.

Previous studies on prokaryotic translation also suggest mRNA structure and accessibility near the translation start site play important roles in translation initiation, a rate limiting step of translation (Studer and Joseph 2006; Gualerzi and Pon 2015; Mustoe et al. 2018). Though we find no significant correlation of the phenotype with change in GC content or number of base changes upon mutations in these regions, we suspect that local mRNA structural changes upon mutations in the 5’ end of the *ccdA* gene affect the translation initiation of CcdA while those in the 3’ end of the *ccdA* gene affect the translation initiation of CcdB, leading to the large codon-specific phenotypic variation observed at both termini of the *ccdA* gene. While the RNA structures of the *ccdAB* transcript predicted by the MFold (Zuker 2003) and RNAfold (Gruber et al. 2008) web servers show that several single codon mutations can affect the predicted mRNA structural architecture of *ccdA* mRNA and its stability, no consistent difference is observed amongst the structures for mutants displaying neutral, inactive and hyperactive phenotypes. Since mRNA stability, transcription and translation are coupled in prokaryotes, to understand mutational effects of mRNA structure on protein levels and activity, one must determine the *in vivo* mRNA structure of the mutants which is beyond the scope of the present study.

## Materials and Methods

### Cloning of WT and saturation mutagenesis library of CcdA

A 986 bp region comprising of the complete *ccdAB* operon along with its upstream and downstream regulatory regions (based on F-plasmid sequence deposited in GenBank) was synthesized at GenScript (USA) and cloned in the EcoRV site of the pUC57 plasmid vector. A few mutations were introduced in this cloned *ccdAB* construct relative to F-plasmid sequence, in order to introduce restriction enzyme sites to facilitate cloning of *ccdA* or *ccdB* mutant libraries (Supplementary Figure S1). A single-site saturation mutagenesis (SSM) library of *ccdA* in this operonic construct was then constructed using inverse PCR methodology(Jain and Varadarajan 2014) (see Supplementary Methods)

### Sample preparation for deep sequencing and data processing

The plasmid library purified from the resistant Top10 gyr strain was considered as the initial unselected library. This was then transformed into the sensitive Top10 strain for selection. We used high efficiency electro-competent cells of Top10 and Top10 gyr strains, having equal efficiencies of 10^9^ CFU/µg of pUC57 plasmid DNA. The detailed method for construction of the Top10 gyr strain is available in Supplementary Methods in the Supplementary Information. Approximately equal numbers of transformants (∼10^8^ in number) from both strains obtained after 14 hours of growth at 37°C on LB-Agar plates containing 100μg/mL Ampicillin,were scraped into 10 mL LB media, and directly used for plasmid purification using a Thermo Scientific Plasmid purification kit for both the unselected and selected libraries. These transformations and subsequent plasmid isolation procedures were performed in three biological replicates. The CcdA gene was PCR amplified from each purified plasmid sample, with primers (Sigma-Aldrich) containing condition specific, six-base long Multiplex Identifier (MID) tags. The resulting 270 bp long PCR products containing the full *ccdA* gene, were pooled, gel-band purified, and sequenced using Illumina Sequencing, on the NovaSeq6000 Platform, at Macrogen, South Korea. The sequencing was performed in a paired-end manner. The overall quality of the sequencing data was assessed using the FASTQC software. Further analysis was performed using our in-house Perl scripts. Reads were separated into ‘bins’ based on their MID tags. The downstream primer sequence was used to identify forward and reverse reads in each bin. A Phred score cutoff of 20 and a minimum read length cutoff of 75 were used to filter low quality reads. Reads were converted into FASTA format and aligned with the *ccdA* gene sequence using the WATER program of the EMBOSS package (Rice et al. 2000; Carver and Bleasby 2003) Default values were used for all the parameters except the Gap Opening Penalty, which was increased to 20. For the present CcdA DMS library, we have used a paired end sequencing platform. Forward and reverse reads of the same read pair were merged together if the overlapping regions were identical and all mismatches between forward and reverse reads were discarded. Only those reads which cover the entire *ccdA* gene length were considered for further analysis. Reads with insertions, deletions and multiple mutations were omitted, and only mapped single mutants were used for further data analyses. The total number of usable reads including the WT reads in the three replicates were 201938, 478296 and 100205. Since NNK primers were used for construction of mutant library, 2.7% of all constructs are expected to be WT sequence. However, deep sequencing of unselected library (as well as Sanger sequencing of 15 colonies) revealed ∼10% of the constructed *ccdA* library to be WT. We suspect the additional 7.3% arises from carry-over of the WT plasmid used as template in mutagenesis PCR reactions, that has not been completely removed by the DpnI digestion and subsequent clean up steps.

We used 200X theoretical coverage for each mutant for deep sequencing and have achieved high average read counts of each mutant in the unselected library (mean = 263 and median=101). The average quality score for this final dataset is 36.76 and majority reads have Q score > 35. Assuming a Q score of 35, the probability of a sequencing error is 1/3162. Assuming that all four bases are equally likely to be assigned in error at a given position, the probability of erroneously observing a specific base substitution at a specific position in a read is 1/12649. For a given error probability and total number of reads, the probability of getting at least the observed number of reads for a specific mutant codon can be calculated assuming a binomial distribution. This binomial cumulative distribution function can be approximated by a Poisson cumulative distribution function. For 100,000 total reads and a probability p<0.05, the minimum number of reads is 13 for a single base substitution when Q35 score is taken. This means that any single–site mutant (with 1 nucleotide change with respect to WT) with 13 or more reads has a probability of <0.05 of being observed by chance. In present analysis, we have discarded all mutants with less than 20 reads in the unselected library. This read cutoff, combined with high average read number ensures the accuracy of the ES_seq_ values and the correctness of phenotypic assignments from the DMS data.

### Assignment of mutant phenotypes based on the deep sequencing data

Read numbers for mutants at all 71 positions (2-72) in CcdA were analyzed. Mutants with <20 reads in the resistant strain were not considered for analysis. The total number of reads in different conditions was calculated. Read numbers for each mutant at a given condition were normalized to the total number of reads in that condition. Mutational enrichment was defined as the ratio of the normalized reads after selection to the normalized reads before selection, for a given mutant. We further normalized the mutant scores with respect to the WT fitness score to obtain the Enrichment score.

Survival score of a mutant i

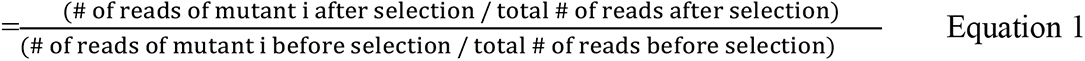

Where total # of reads refers to all single mutants and WT reads identified in a sample.

Enrichment score of a mutant i, 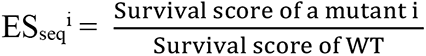

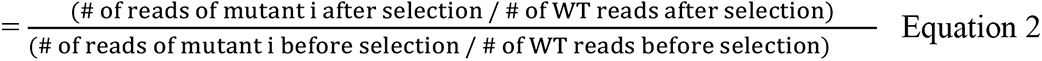

ES_seq_ is a measure of the enrichment (growth advantage) of a mutant relative to the WT construct in the sensitive strain. The ES_seq_ scores were calculated for all mutants in the library that had >20 reads in the resistant strain, separately for each replicate. ES_seq_ values used in the work are the average of the three replicates.

For each residue in the CcdA protein, reads for all 32 possible mutant codons (NNK codons) were analyzed in the sensitive vs. resistant strain, and phenotypes were assigned in terms of the ES_seq_ scores. Positions showing codon specific mutation effects were analyzed separately.

### Fitness (w) estimation

Fitness was calculated as previously described (Bailey et al. 2014; Lebeuf-Taylor et al. 2019) using the formula,

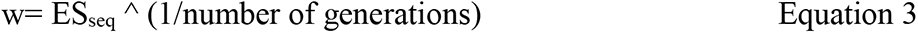

The number of generations was taken as 37 and calculated using a lag time of 1.4 hours and doubling time of 20.2 minutes as reported for *E. coli* growth on plates (Fujikawa and Morozumi 2005).

### *In vivo* activity of individual single-site CcdA mutants

Selected single-site synonymous mutants of CcdA (in pUC57-*ccdAB* construct) were designed and synthesized from GenScript and sequence confirmed (see Table 1). 40 ng of each plasmid was individually transformed into the sensitive and the resistant strains and plated on LB-amp agar plates in ten-fold serial dilutions and grown at 37°C for 16 hrs. The number of CFUs for the transformants in Top10 strain obtained in different dilution plates was counted to confirm their *in vivo* activity. For easy readout, serially diluted transformation mix samples were also spotted using 3μL volumes per spot to visualize differences among mutant phenotypes.

### Monitoring the relative levels of CcdA and CcdB using a quantitative proteomics approach

The relative levels of CcdA and CcdB proteins *in vivo* were quantified from cell lysates using mass spectrometry based quantitative proteomics. All the experiments were carried out in two or more technical replicates except for F plasmid sample. Detailed Methodology is available in the Supplementary Methods section. The ratio of CcdA: CcdB protein levels were determined for each replicate and the mean values with standard deviations have been reported.

### Estimation of *ccd* specific mRNA levels using qRT-PCR

For RNA isolation and quantification, 3 ml of culture was grown for each of the mutants of *ccdA* transformed in *E. coli* Top10 gyr strain. Cells were grown to saturation under shaking conditions at 37°C, 180rpm, pelleted and total RNA was extracted by the RNAsnap method (Stead et al. 2012). Chromosomal and plasmidic DNA was removed by treatment with 2 units of DNase1 (New England Biolabs) for 2 hours at 37°C, followed by Sodium Acetate-ethanol precipitation of RNA. RNA was quantified by nanodrop spectrophotometric estimation, and quality was assessed by agarose gel electrophoresis (samples with A260/A280 = 2, were used for further studies), prior to downstream processing. 2µg of total RNA was taken in a sterile, RNase-free microcentrifuge tube and 0.5µg of random hexamers (Sigma-Aldrich) was added to serve as primer for cDNA synthesis along with DEPC (Sigma-Aldrich) treated double autoclaved MilliQ water. The mixture was heated to 60°C for 10 minutes to melt secondary structure within the template. The tube was cooled immediately on ice to prevent secondary structure from reforming. To this mix, 200 units of SuperScript III Reverse Transcriptase (Invitrogen) were added, along with dNTPs, RT Reaction Buffer and 25 units of Ribonuclease Inhibitor following the manufacture’s protocol. The reaction mix was incubated for 60 minutes at 37°C. cDNA was directly used for quantitative PCR (Q-PCR) analysis. Q-PCR was set up with *ccdA* as well as *ccdB* gene specific primers (250nM) using the cDNA template and Bio-Rad iQ SYBR Green Supermix (1x) in 20 µl total reaction volume, in a Bio-Rad iQ5 machine. The thermocycle parameters used for the Q-PCR were initial 95°C for 10 mins followed by 95°C for 30 secs, 56 °C for 30secs and 72°C for 30 secs in 40 repeat cycles. 16S rRNA was used as the internal positive control for total RNA quantification. Reactions with no reverse transcriptase as well as no template were used as a negative control.

Threshold cycle of amplification in PCR reaction was analyzed and automatically provided by the Bio-Rad iQ5 Optical System Software Version 2.1. All reactions were carried out in triplicates. The negative control experiments had Ct values in the range of 29-31.

### Computational prediction of mRNA secondary structure

The initial 150 bp of the *ccdAB* transcript were submitted to mFOLD (Zuker 2003) and RNAfold (Gruber et al. 2008) for prediction of secondary structure. All energy parameters were set at default values. Detailed output was obtained in the form of structure plots with reliability information, single strand frequency plots, and energy dot plots. Local mRNA secondary structural elements encompassing the region of synonymous inactive mutations were further analyzed.

### Calculation of parameters describing various DNA sequence features

The codon usage frequency of *E.coli* K12 strain (Nakamura et al. 2000)(based on 14 coding gene sequences) was used in the study since the Top10 strain is a derivative of the K12 strain. These values of codon usage frequency in K12 was also found to correlate very well (r = 0.98) with the*E. coli*genome codon usage frequency table at the GenScript website (https://www.genscript.com/tools/codon-frequency-table). The codon usage frequency fraction (CUF) was used to calculate the relative codon usage (RCU) for mutants, using the following formula:

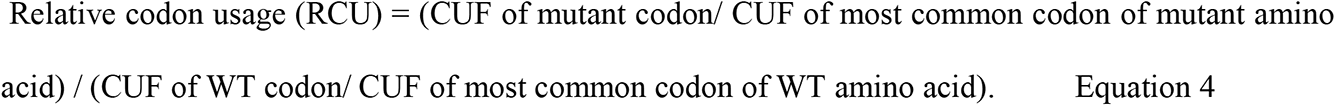

The Codon Adaptation Index, CAI for WT CcdA sequence and all single-site mutants were calculated using a Python implementation of CAI and 274 highly expressed genes from *E.coli* K12 as reference (Lee 2018).

The tRNA abundance raw data (Dong et al. 1996) depicts the abundance of tRNA molecules in an *E. coli* cell that decode one or more codons. We assigned the tRNA abundance values to the respective codons that they decode, and calculated the relative tRNA abundance values associated with each codon using a similar normalization as described above for calculation of relative codon usage.

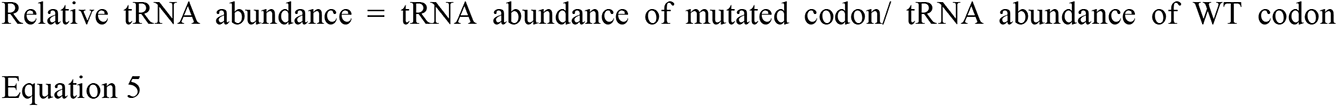

The ribosome stalling data (Chevance et al. 2014) describes the observed averaged pausing of ribosome on each codon at the A-site for all genes in *E. coli*. The ribosomal pausing was estimated by introducing each codon individually into a 6X-His leader sequence of the histidine operonic system and quantified by downstream HisD-LacZ fusion protein (β-lactamase) activity. For the current work, the relative ribosome stalling for each mutation was calculated by dividing the mutant codon pause value by the WT codon pause value.

The relative GC content for mutations was calculated by dividing the fractional GC content of mutant codon by that of WT codon. The number of base changes is the number of positions in a 3 bp codon where the mutant codon varies from the WT codon.

To investigate if there is any significant difference in the distribution of these sequence parameters across the inactive and active classes of mutants, we chose to use the non-parametric Mann-Whitney test since our data is ordinal and does not follow a normal distribution.

### Calculation of anti-Shine Dalgarno interaction energies for synonymous mutants in *ccd* mRNA

The difference in the interaction energy between single synonymous mutants in *ccd*A and the WT sequence, with the consensus anti-Shine Dalgarno (aSD) sequence (5’ CCUCCUAU 3’), was calculated for a window of eight nucleotides using the RNAsubopt program from the RNA Vienna package 2.4.3 (Gruber et al. 2008). The calculations were performed for ten such windows encompassing the mutant codon. The average energy difference for each mutant across these ten windows was compared for all the available synonymous active and inactive mutants. Such aSD like sequences are known ribosomal pause sites (Li et al. 2012).

The binding energies for different mutants at the ribosome binding site of CcdB in the *ccdA* gene (residue positions 70 and 71) to the anti-Shine Dalgarno (aSD) sequence (5’ CCUCCUAU 3’) was also calculated similarly, to look for possible *ccdA* mutants (at residue positions 70 and 71) that can alter the translation initiation of CcdB.

### Yeast surface display of CcdA library and single mutants to probe CcdB binding

The CcdA libraries, individually recovered both from the resistant and the sensitive strains, were amplified from the pUC57-*ccdAB* plasmid vector and recombined into the yeast surface display vector, pETcon (Addgene plasmid # 41522) between NdeI and XhoI sites, using yeast homologous recombination in *S. cerevisiae* EBY100 strain. Positive clones, sufficient in number to cover the library diversity, were obtained on selective SDCA plates. The transformant pools obtained were grown in liquid SDCAA media for 48 hrs and stored in aliquots of 10^9^ cells per ml in SDCA media containing 20% glycerol at −70°C. The same methodology was used for cloning WT and individual synonymous mutants of CcdA into pETcon. The CcdA coding region was cloned as a C-terminal fusion to the yeast Aga2p protein with a C-terminal c-myc tag under control of the GAL promoter. The library was displayed on the yeast surface as described (Chao et al. 2006). Cells were induced and grown in SGCA media containing 2% galactose at 30°C for 16 hrs. The surface expression of CcdA was monitored by incubation of 10^6^cells with anti-c-myc antibody raised in chicken (1:400 dilution), that binds to the Myc tag at the C-terminus of CcdA. Anti-chicken IgG conjugated AlexaFluor-488 (1:300) was used as the secondary antibody. CcdB binding to the surface expressed CcdA was assessed by incubating cells with a fixed concentration of 2nM biotinylated CcdB (or varying concentrations between 0.1pM and 200nM for titration experiments) and binding of streptavidin conjugated AlexaFluor-633 secondary antibody (1:2000) to biotinylated CcdB was monitored. All the primary and secondary antibodies were obtained from Invitrogen. Labelled cells were analyzed on a BD FACSARIA III. The titration curves were fit to a single ligand binding model to obtain the dissociation constants as described previously (Rathore et al. 2017; Chandra et al. 2021). Double plots for mean fluorescence intensities for both expression and binding for the CcdA library as well as for single synonymous mutants were analyzed to monitor differences with respect to the WT.

### Monitoring the effect of single synonymous mutations in the RelBE operon

#### Design of mutants

The WT sequence for the RelBE operon from *E.coli* K12MG1655 strain was retrieved from the NCBI database. The entire transcription unit of 910 bp (Bech et al. 1985) including another gene *relF*, which is expressed as a part of the same operon, was cloned in the EcoRV site of the pUC57 vector at GenScript. Five different constructs having synonymous mutations in the *relB* gene were synthesized with a stop codon in the *relE* toxin gene to ensure normal cell growth even after inactivation of the antitoxin, for ease of cloning. Three base synonymous substitutions are only possible with serine codons. Three base synonymous substitutions with varying codon preferences were made in three of the serine codons, S3, S16 and S28, all at the N-terminal half of the *relB* coding sequence. One representative two base substitution at position R7 was also tested. A single base substitution at the D77 residue was made to increase the strength of the SD sequence for *relE* and was used as a positive control mutant. To obtain an easy phenotypic readout, the introduced stop codon in the *relE* gene was reverted, using the inverse PCR strategy described earlier (Jain and Varadarajan 2014).

#### Phenotype testing

Due to absence of a strain that is resistant to toxin action, we employed a toxin deletion strain (*E. coli* BW25113 Δ*relE*) for the propagation of mutants. By virtue of this deletion, the WT chromosomal operon is de-repressed, leading to an excess of antitoxin in the cell, relative to the WT strain (*E. coli* BW25113 *relBE*). The inverse PCR products for individual mutants were phosphorylated and ligated, as described previously. Ligation mixes were individually transformed into the *E. coli* BW25113 ΔrelE strain, to recover the plasmid. Single clones were isolated, and sequence confirmed. Equal amounts of the mutant plasmids (∼30 ng) were transformed in *E. coli* BW25113 WT strain (having chromosomal *relBE* genes intact) to test their phenotype, transformation mixes were plated on LB-amp agar plates in ten-fold serial dilutions and grown at 37°C for 16 hrs. The number of CFUs for the transformants in *E. coli* BW25113 WT strain obtained in different dilution plates was counted. Serially diluted transformation mix samples were also spotted using 2μL volumes per spot to visualize differences among mutant phenotypes.

## Author Contributions

K.G. prepared the SSM library. S.C. carried out the experiments except for the quantitative proteomics (S.V.M. and H.G.) and the experiments with RelBE system (P.K). S.K. processed the deep sequencing data. A.A helped in acquiring FACS data. R.V, S.C. and K.G. designed the experiments, analyzed the overall data and wrote the manuscript with critical inputs and review from all other authors.

## Data and Material availability

The data relevant to the figures in the paper have been made available within the article and in the Supplementary Information section, or are submitted to GitHub. The deep sequencing data are submitted in NCBI SRA database and to be released upon acceptance of manuscript (BioProject ID: PRJNA795874). The deep sequencing read counts, calculated Fitness Scores as well as values for codon usage, tRNA abundance, GC content, number of base changes, ribosome stalling, predicted binding energies for DNA interactions with SD and aSD sequences and all data generated in experiments such as YSD, Q-PCR, proteomics are available in GitHub (https://github.com/rvaradarajanlab/CcdA-Operonic-Data.git). The raw quantitative proteomics data is available at https://db.systemsbiology.net/sbeams/cgi/PeptideAtlas/PASS_View?identifier=PASS01727. All materials generated in this study are available from the lead contact without restriction.

## Acknowledgement

SC acknowledges Ministry of Human Resource Development for her fellowships and thanks all the members of the RV lab for their valuable suggestions. We thank Professors SP Arun and Mrinal Ghosh, IISc for their valuable inputs on error probability calculations for deep sequencing reads and Munmun Bhasin for her help with the calculations, and for useful discussions. We also acknowledge funding for infrastructural support from the following programs of the Government of India: DST FIST, UGC Centre for Advanced study, Ministry of Human Resource Development (MHRD), and the DBT IISc Partnership Program.

