## Supplementary Information for "The high mutational sensitivity of *ccdA* antitoxin is linked to codon optimality"

#### Supplementary Figures

GGAGATGGCCAAAACCCCAAGTTACGGATCTTCCTCTCCCTCCGCACAGCGTTACATCCCGTCAGCACAGCATGT

AGTGCCTCATACAGTTGCCCATGGCACTATATGTTGTGTTGTATCTCTGGACTGTGATGCGCCGCGCAGGGGCGG

Eco RI

AAAACAGCGATATGATGATTTTCTCAGCGTTGTACACTTCCGGAAAGTCGTTTATTCAAATAAAGTCGAATTCATAC

GAAACGGGAATGCGGTAATTACGCTTTGTTTTTATAAGTCAGATTTTAATTTTTATTGGTTAACATAACGAAAGGTA

PsiI

AAATACATAAGGCTTATAAAAAGCCAGATAACAGTATGCGTATTTGCGCGCTGATTTTTGCGGTATAAGAATATATAC

Bgl 2

TGATATGTATACCCGAAGTATGTCAAAAAGATCTGTGCTATGAAGCAGCGTATTACAGTGACAGTTGACAGCGACA

GCTATCAGTTGCTCAAGGCGTATGATGTCAATATCTCCGGTCTGGTAAGCACAACCATGCAGAATGAAGCCCGTCG

TCTGCGTGCCGAACGCTGGAAAGCGGAAAATCAGGAAGGGATGGCTGAGGTCGCCCGGTTTATTGAAATGAACG

Dra I

GCTCTTTTGCTGACGAGAACAGGGACTGGTGAAATGCAGTTTAAAGTTTACACCTATAAAAGAGAGAGCCGTTATC

GTCTGTTTGTGGATGTACAGAGTGATATTATTGACACGCCCGGGCGACGGATGGTGATCCCCCTGGCCAGTGCAC

GTCTGCTGTCAGATAAAGTCTCCCGTGAACTTTACCCGGTGGTGCATATCGGGGATGAAAGCTGGCGCATGATGA

CCACCGATATGGCCAGTGTGCCGGTCTCCGTTATCGGGGAAGAAGTGGCTGATCTCAGCCACCGCGAAAATGACA

Bam HI

TCAAAAACGCCATTAACCTGATGTTCTGGGGAATATAAATGTGAGGATCCGTTATACAC

**Figure S1. Sequence of *ccdAB* operon used in the study.** Sequence of *ccd* operon synthesized and cloned in pUC57 vector at GenScript (USA). Restriction sites have been underlined and marked. The *ccdA* coding region is in green and *ccdB* is in purple. Mutations in the construct with respect to the native F-plasmid borne *ccd* sequence were introduced to facilitate cloning of the *ccdA* and *ccdB* libraries and are shown in red. Following the *ccdB* gene, the first few residues of the *resD* gene have been included with a stop codon at the second residue position and a mutation to introduce a BamH1 site.

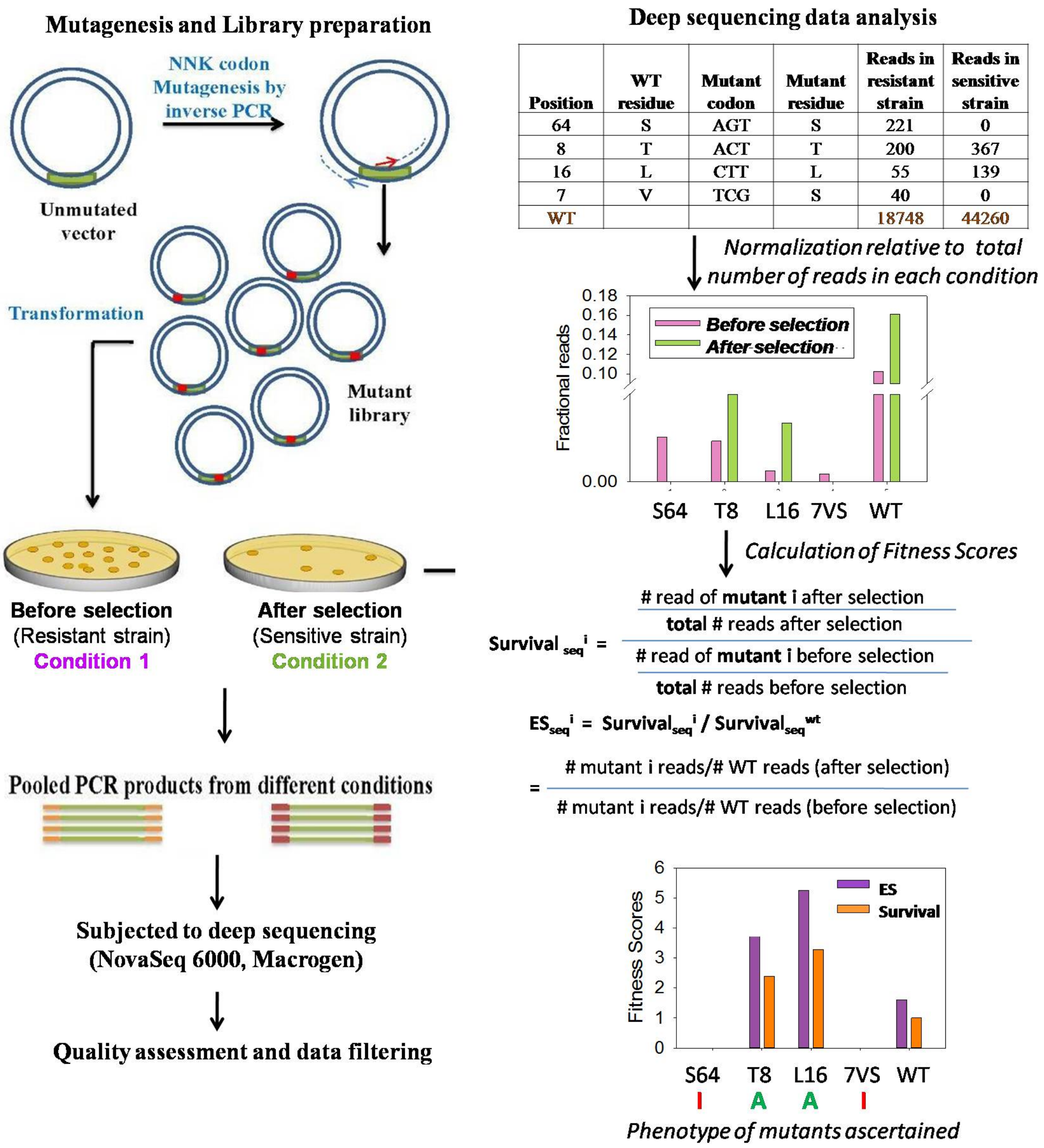


**Figure S2. Saturation mutagenesis coupled to deep sequencing to infer mutational sensitivities.** Single-site saturation mutagenesis libraries are generated using inverse PCR based methodology that involves use of non-overlapping primers with the mutant NNK codon at the 5’ end of the forward primer. Mutant phenotypes are assayed by transforming into resistant (condition 1) and sensitive strains (condition 2). Pooled plasmids from different conditions are used as template to amplify the gene of interest with primers having condition specific tags. Pooled PCR products are subject to deep sequencing (NovaSeq6000 platform, Macrogen) in three biological replicates and the data is analyzed to obtain the relative enrichment scores ($ES_{seq}$) of the mutants relative to WT (*See Methods*). Enrichment Score ($ES_{seq}$) of mutant is calculated using the fraction of reads for the mutant after selection (condition 2) to that before selection (condition1), relative to WT. The $ES_{seq}$ is therefore the survival score of the mutant normalized with respect to WT. Thus for WT, $ES_{seq}$ =1. Mutants with lower values of $ES_{seq}$ show reduced growth in the sensitive strain, relative to WT. Mutants with $ES_{seq}$ $\leq$ 0.1 are classified as inactive (I), while mutants with $ES_{seq}$ scores > 0.1 classified as active (A).

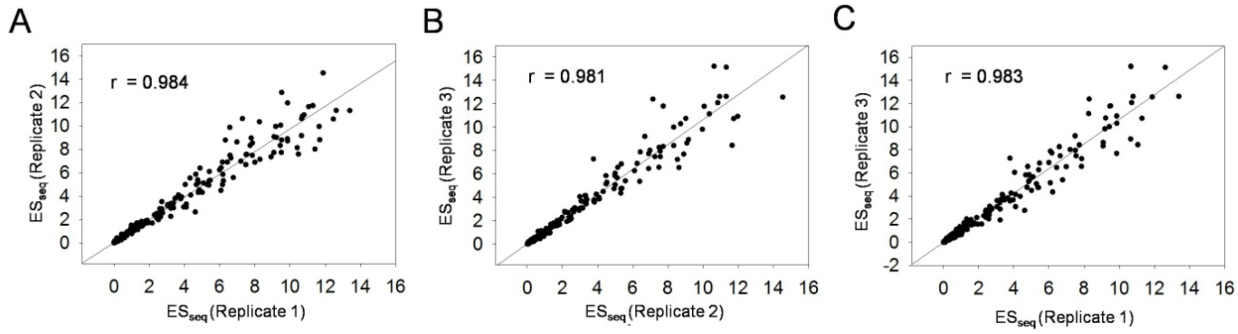


**Figure S3. The phenotypic scores ($ES_{seq}$) of mutants are highly correlated between biological replicates**

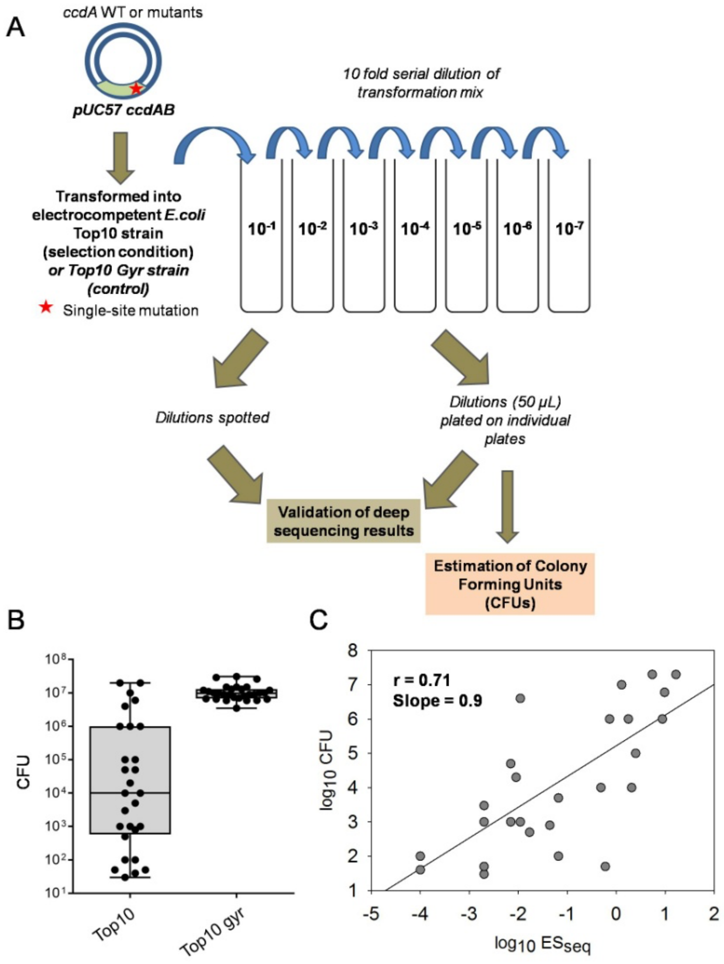


**Figure S4. Validation of deep sequencing results in individual mutants of CcdA.v**(A) Transformation of Top10 *E.coli* (CcdB-sensitive) strain with WT and individual CcdA mutants cloned in pUC57 plasmid vector as part of the *ccdAB* operon, followed by serial dilution and phenotypic assay. 50µl of serial dilutions $10^{-1}$ to $10^{-7}$ of transformation mix of all mutants were plated on individual LB-amp agar plates. The number of colonies was counted for different dilutions after overnight growth at 37°C and were used to calculate the CFU/ml (averaged over results of the two highest dilution with countable colonies) for each construct. All constructs were also transformed to Top10 gyr *E.coli* (CcdB-resistant) strain that served as transformation control. (B) The distribution of CFU observed for the different *ccdA* mutants after transformation in the Top10 and Top10 gyr strains. While the numbers of CFU were near identical in Top10 gyr strain, we observed large variability in cell survival in the Top10 strain across the different *ccdA* mutants. (C) Correlation of the calculated CFU values for different *ccdA* mutants with corresponding $ES_{seq}$ values obtained from deep sequencing results. Plate and deep sequencing data are well correlated.

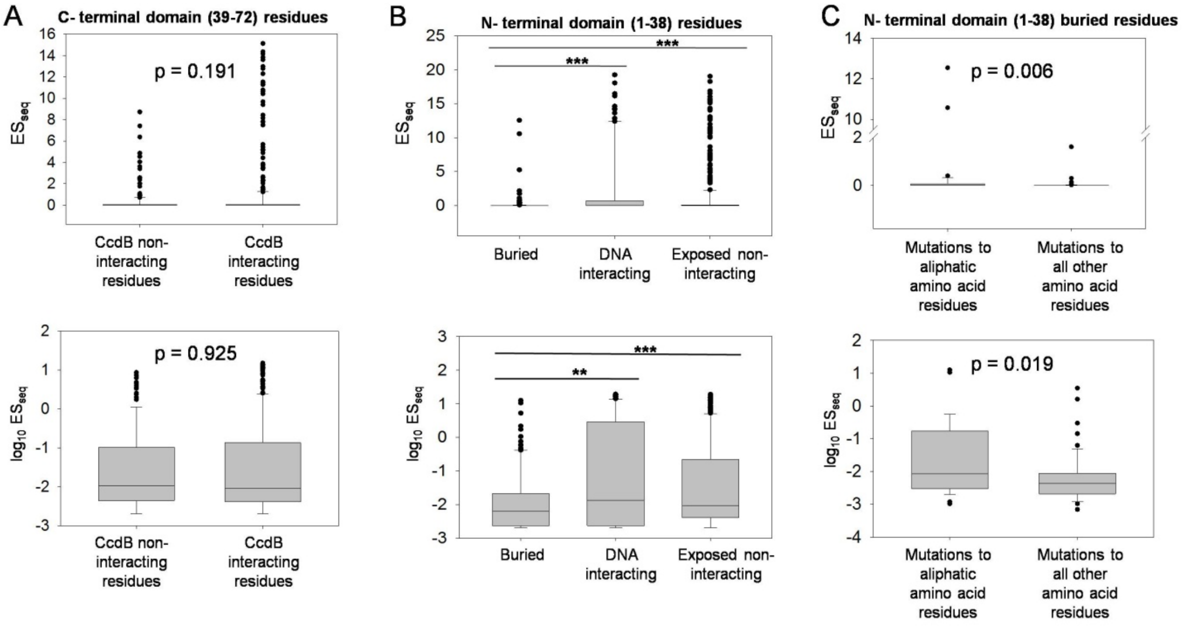


**Figure S5. Buried residues in the structured N-terminal domain have higher mutational sensitivity than all other residues.** Distribution of $ES_{seq}$ values for all *ccdA* mutants investigated by deep sequencing was studied to understand the role of different classes of functional residues. (A) Box plots of distribution of $ES_{seq}$ values in absolute (top) and logarithmic scale (bottom) for CcdB interacting and CcdB non-interacting residues in the disordered C-terminal domain, classified based on the available structure of CcdB bound to the CcdA C-terminal domain (PDB ID : 3G7Z). There was no statistically significant difference between phenotypes of mutations at CcdB interacting and non-interacting sites. (B) Box plots of distribution of $ES_{seq}$ values in absolute scale (top) and logarithmic scale (bottom) for different N-terminal domain residue classes. The classification of residues was done based on the available structure of the dimeric N-terminal domain of CcdA bound to DNA of the operator/promoter region of the *ccdAB* operon (PDB ID : 2H3C). Mutations at buried residues have considerably lower $ES_{seq}$ values (*** means p value < 0.001 and ** means p value < 0.01) than mutations at any other class of residues. (C) Box plots of distribution of $ES_{seq}$ values in absolute (top) and logarithmic scale (bottom) for the aliphatic (A, C, L, I, M and V) and all other amino acid substitutions in the N-terminal domain buried residues. The substitutions to all non-aliphatic residues, at the N-terminal domain buried positions show significantly lower $ES_{seq}$values than substitutions to aliphatic residues, leading to more drastic inactive phenotype. Statistical significance was assessed by a Mann-Whitney Rank Sum Test. A p-value > 0.05 indicates that there is no statistically significant difference between the distributions of the indicated parameter for active and inactive mutants, at the 95% confidence interval.

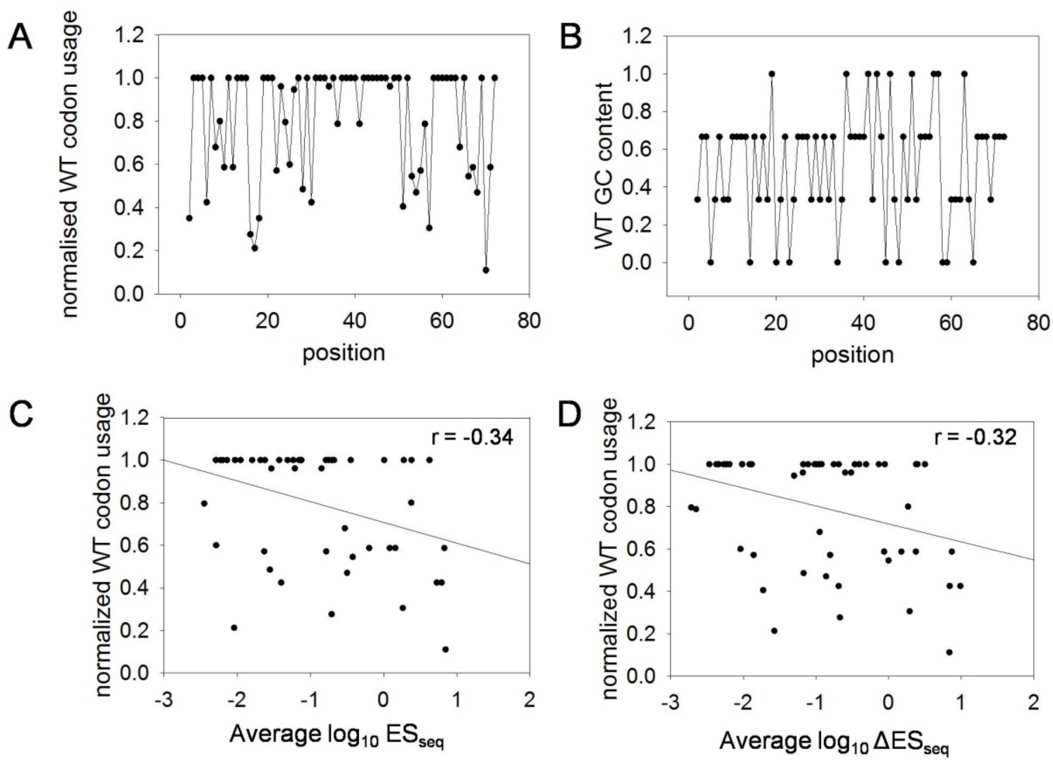


**Figure S6. Unequal distribution of inactive phenotypes of mutants across gene length can be partially explained by the non-uniform distribution of rare and preferred codons in the *ccdA* gene.** (A) Normalized WT codon usage was calculated by dividing the *E. coli* codon usage of the WT codon by codon usage of the most common codon. A WT codon with normalized codon usage value of 1indicatesthat the WT codon is the most optimal or commonly used in the *E. coli* genome. Interestingly, the central region of CcdA has high numbers of optimal codons, while rarer codons populate the extremities of the *ccdA* gene. (B) GC content of WT codons of the CcdA gene as a function of residue positions. (C) Normalized codon usage values for each residue in the WT *ccdA* gene show modest negative correlation with corresponding position average $log_{10}ES_{seq}$. This indicates that residue positions bearing optimal WT codons are more likely to produce inactive phenotypes upon mutations. (D) Normalized codon usage values for residue positions in WT *ccdA* gene also show a modest negative correlation with average $log_{10}\Delta ES_{seq}^{codon}$ values (calculated as described in Figure 2B), indicating that there is larger variation in phenotypes among degenerate codons at residues positions bearing non-optimal WT codons.

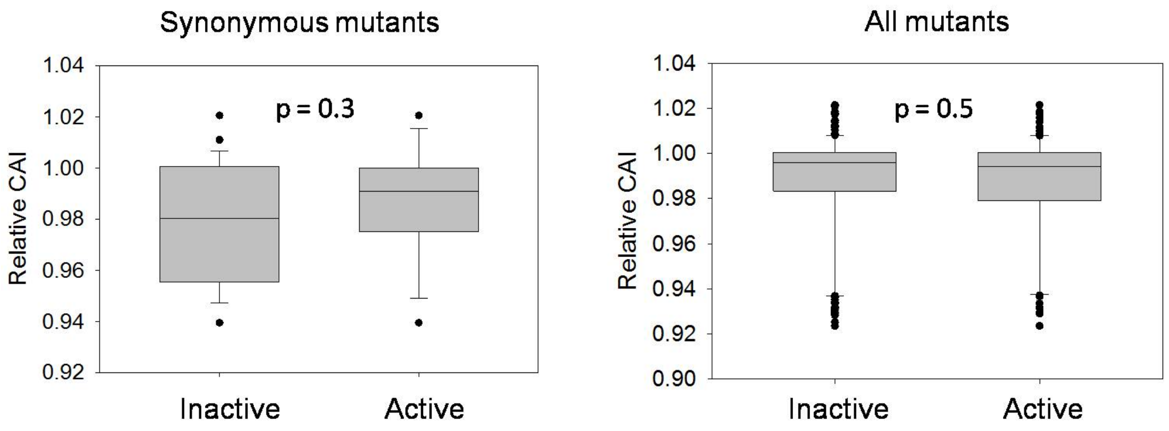


**Figure S7. Distributions of Relative Codon Adaptation Index (CAI) for inactive and active *ccdA* synonymous (left panel) and all (right panel) mutants.** Relative CAI = CAI calculated for mutant *ccdA* sequence/CAI of WT *ccdA* sequence. Statistical significance was assessed by a Mann-Whitney Rank Sum Test.

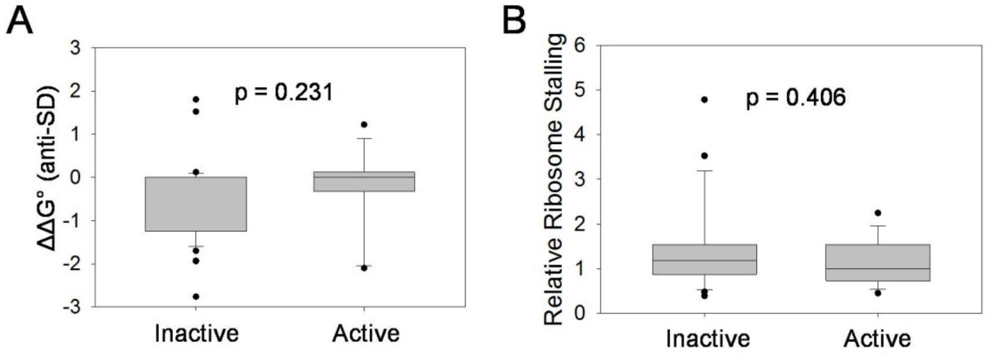


**Figure S8. Effects of potential ribosomal pause sites on phenotype of CcdA synonymous mutations.** (A) Distribution of differences in anti-SD sequence interaction energies (in kcal) amongst inactive and active synonymous mutants. The change in binding energy to the anti-Shine Dalgarno sequence on the ribosome in a mutant with respect to the WT sequence was calculated for a window of 10 bases using the RNAsubopt program in the Vienna RNA package. (B) Distribution of codon specific relative ribosome stalling (Chevance et al. 2014) amongst inactive and active synonymous mutants. Statistical significance was assessed by a Mann-Whitney Rank Sum Test. A p-value > 0.05 indicates that there is no statistically significant difference between the distributions of the indicated parameter for active and inactive mutants, at the 95% confidence interval.

### Supplementary Methods

##### Preparation of a single-site saturation mutagenesis library of CcdA

Mutagenic primers for all 71 positions (2 - 72) of CcdA were designed such that the degenerate codon (NNK) was at the 5' end of each 21 bp forward primer. A non- overlapping adjacent 21 bp reverse primer, along with the forward primer, was used to amplify the entire pUC57-*ccdAB* plasmid by inverse PCR methodology using Phusion DNA polymerase (Jain and Varadarajan 2014). The primers were obtained in 96-well format from the PAN Oligo facility at Stanford University. A master-mix was made for performing the PCR reactions for all positions in a 96 well format. Following quantification, an equal amount of PCR product (~200ng) at each position was added to get a pool of PCR products of all required fragments. NEB DpnI treatment and gel-band purification of the pooled PCR product at the required size (~3.6 Kb) was done using a Fermentas GeneJET Gel Extraction Kit. After purification, pooled PCR product was phosphorylated, followed by ligation. The ligated product was transformed into high efficiency ($10^9$ CFU/µg of pUC57 plasmid DNA) electro-competent *E. coli* Top10 gyr cells. 5µg of the total ligated PCR product was transformed into 2mL of competent cells (4 aliquots of 500µl each) and the cells were plated on LB agar plates containing 100µg/mL ampicillin, for selection of transformants. Plates were incubated for 18-20 hrs at 37˚C. Around 2000-3000 colonies obtained on each plate were washed off from a total of 8 LB agar plates (150mm diameter) into a total volume of 100 mL of LB media. Pooled plasmid library was purified using a Qiagen plasmid maxiprepkit and is treated as the initial complete *ccdA* library.

##### Construction of Top10 gyr strain

The *E. coli* Top10 gyr strain was constructed using homologous recombination-based methodology (Datsenko and Wanner 2000). Briefly, a GyraseA fragment of DNA Gyrase enzyme having the mutation R462C was amplified from the parent strain *E. coli* CSH501. To enable insertion of this fragment at a particular locus in the *E. coli* Top10 genome, the fragment had homology at the 5` end with the Gyrase sequence upstream of the mutation and at the 3` end with *kan* gene from pKD4 plasmid. The Kan cassette was added for selection of recombinants. The *kan* gene containing FRT sites, was amplified such that it has homology at the 3' end with Gyrase sequence, downstream of the R462C mutation. This Gyrase-kan cassette was transformed in *E. coli* Top10 strain by electroporation and transformants growing on Kanamycin plates were selected. The mutation was confirmed by DNA sequencing. The *kan* gene was later removed by transforming the strain with plasmid pCP20 expressing the FRT-FLP recombinase, as described previously (Datsenko and Wanner 2000).

##### Primers used for qRT-PCR

The sequences for the forward and reverse *ccdA* specific primers were

5’GAAGCAGCGTATTACAGTGACAGTTG3’ (binding 2-28 bp region of *ccdA*) and 5’CGTCAGCAAAAGAGCCGTTCA3’ (binding 181-202 bp region of *ccdA*) respectively.

The sequences for the forward and reverse *ccdB* specific primers were 5’GAGAGAGCCGTTATCGTCTGTTTGTG3’ (binding 28-54 bp region of *ccdB*) and

5’GATAACGGAGACCGGCACAC3’ (binding 208-228 bp region of *ccdB*) respectively.

Both the gene specific amplicons for the Q-PCR reaction were 200bp.

The sequences for the forward and reverse 16S rRNA specific primers were

5’ CCTAACACATGCAAGTCGAACGG 3’ and

5’ CTCCATCAGGCAGTTTCCCAGAC 3’ respectively.

**Monitoring the relative levels of CcdA and CcdB using a quantitative proteomics approach**

1. ***Selection of proteolytic peptides from CcdA and CcdB for MRM experiments:***

In order to identify proteolytic peptides, *E. coli* lysates were resolved on SDS-PAGE, and gel bands corresponding to CcdA and CcdB were excised and subjected to proteolytic digestion using trypsin. Peptide digests were analyzed on a Thermo Orbitrap Fusion$^{TM}$ Tribrid$^{TM}$ Mass Spectrometer interfaced with Easy-nLC 1000 nanoflow liquid chromatography system (Thermo Scientific, Bremen, Germany). Peptides were loaded onto a trap column (75 µm x 2 cm, Magic C18AQ, 5 µm, 100 Å, Michrom Biosciences Inc., Auburn, CA) using 0.1% formic acid at a flow rate of 3 µl/min. The peptides were then resolved on an analytical column (75 µm x 20 cm, Magic C18AQ, 3 µm, 100 Å, Michrom Biosciences Inc, Auburn, CA) at a flow rate of 350 nl/min using a linear gradient of 5-35% B (0.1% formic acid in 95% acetonitrile) over 18 min and a run time of 30 min. The MS and MS/MS scans were acquired at a mass resolution of 120,000 and 30,000 at 200 m/z, respectively. Full MS scans were acquired in the m/z range of 350-1550. Precursor ions with single charge or unassigned charge were rejected. Dynamic exclusion of fragmented precursor ions was set to 30 sec. Precursor ions were isolated using a quadrupole mass filter with isolation width of 2 m/z. Higher energy collision dissociation was used as the fragmentation method with 32% normalized collision energy. Proteome discoverer (version 1.4) software suit (Thermo Fisher Scientific, Bremen Germany) was used to analyze the data. Three peptides were selected for CcdA and two for CcdB. Peptides ITVTVDSDSYQLLK and LFVDVQSDIIDTPGR was used for quantitation of protein CcdA and CcdB, respectively while others were used for identification purposes only.

2. ***Mass spectrometry protein sample preparation:***

To quantify the levels of CcdA and CcdB proteins in the *E.coli* lysate, 10 ml cultures of *E. coli* Top10 gyr cells transformed individually with different CcdA mutants were grown till saturation (OD600 of

1.0) under shaking conditions at 37˚C, 180rpm. Cells were pelleted, washed twice with PBS, and lysed by sonicating in lysis buffer (2% SDS in 100mM triethyl ammonium bicarbonate (TEABC) buffer, pH 8.5). The lysates were clarified by centrifugation at 12,000 g for 30 min at 4°C. Proteins were precipitated with ice-cold acetone to remove SDS and re-constituted in 100mM TEABC. Bicinchoninic acid (BCA) assay kit (Thermo-Scientific) was used to estimate protein concentration. An equal amount of protein from each sample was reduced with 10mM DTT at 60°C for half an hour. The samples were cooled to room temperature and proteins were alkylated by adding iodoacetamide at a final concentration of 20mM and incubating at room temperature in the dark for 15 min. Proteins were subjected to proteolytic digestion overnight by adding sequencing grade trypsin (Promega) at an enzyme: substrate ratio of 1:20. For LC-MS-MRM experiments, samples were spiked with stable isotope labeled peptides (SpikeTide$^{TM}$ TQL peptides were purchased from JPT Peptide Technologies, Germany) before adding trypsin. After overnight incubation, the reaction was quenched with 10% formic acid and centrifuged at 12,000 g to remove any precipitate. Sample clean-up was performed using reverse phase spin columns (Harvard apparatus) according to the manufacturer's protocol.

##### 3. *SRM/MRM analysis:*

LC-MS/MRM analysis was performed with an electrosprayionization-triple quadrupole mass spectrometer (QTRAP 6500; SCIEX) coupled with an HPLC system (Agilent 1260 systems; Agilent Technologies, Santa Clara, CA). The peptides were resolved on a reverse phase C18 column (Agilent Poroshell 120 EC-C18 2.7µm, 4.6x50mm) at a flow rate of 350 µl/min and a run time of 10 min. The LC method used was as follows: 10% - 30% solvent B from 0-3 min, 30%-50% solvent B from 3-5 min, 50%-95% solvent B from 5-7 min and 95%-5% solvent B from 7-10 min. The data was acquired in SRM/MRM mode using a dwell time of 25 msec. Each target peptide and corresponding SIL peptide were measured. Standard curves were generated by spiking different amounts of SIL peptides (0.5, 0.1, 0.25. 0.5, 1, 5, 10, 20 pmol) into a constant amount of *E. coli* digest background.

##### 4. *LC-MS/MRM data analysis:*

MRM data was analyzed using Skyline 3.5 (MacLean et al. 2010) and composite peak AUCs were calculated after Savitsky-Golay smoothening. Each transition was inspected and XICs for each transition was generated. Response curves were generated by plotting the ratio of heavy: light peptide over different concentrations of spiked heavy peptide. Linear regression analysis was carried out to determine the linearity of the response curve. Also, to validate the analytical method, regression analysis of variance (ANOVA) of the linear regression data measurements (at a 95% confidence level)was performed. In addition, to illustrate the reliability of LC-MS/MRM analysis, the %RSD was calculated, which was within 20% in all cases. Peaks corresponding to endogenous target peptides were identified based on the retention time of the corresponding SIL peptide as it co-elutes with the endogenous peptide. Welch's test was used to determine the statistical significance of abundance differences between the wild type and mutants.
